# Flexible Steering and Conflict Resolution: Pro-Goal/Anti-Goal Gating in *Drosophila* Lateral Accessory Lobes

**DOI:** 10.1101/2025.10.19.683286

**Authors:** Chen-Chieh Liao, Ning Chang, Yi-Shiuan Liu, Chung-Chuan Lo

## Abstract

Navigation comprises multiple processes: sensory integration, position estimation, decision, and finally, motor command generation, which is the control problem. Previous studies discovered that PFL3 neurons compute a heading-goal error and directly project to the descending neurons that generate the steering commands. Yet, this direct pathway is necessary but not sufficient for the flexible control required to pursue goal while avoiding threats and resolving conflicts. Using the Drosophila connectome and circuit modeling, we uncover a layered control architecture in the lateral accessory lobes (LAL) that arbitrates between goal pursuit and stimulus-driving overrides. First, a dorsal pathway that forms a dominant indirect “pro-goal” push-pull circuit that amplifies left-right asymmetries. Second, a faster ventral pathway forms an “anti-goal” circuit that recruits and inverts the pro-goal circuit to redirect movement, and only engages when a goal is present. The architecture also resolves two symmetry challenges: Rear Stalemate (goal 180 behind) and Front Stalemate (threat dead ahead), by rapidly amplifying tiny perturbations into decisive turns. Together, these motifs instantiate comparator, override, and gating principles in a compact neural controller, yielding testable predictions for insect motor control and design rules for bio-inspired, embodied intelligence.

## Introduction

Navigating through complex and dynamic environments is a fundamental cognitive ability that supports survival across the animal kingdom ^1–3^. To reach desired locations and avoid threats, an animal must integrate multiple sources of information: it must estimate its own position and heading ^4–7^, construct internal maps of its environment ^8^, identify and represent behavioral goals ^9–11^ and ultimately generate motor commands to steer its body toward those goals^12,13^. Therefore, navigation is not merely about computing a destination, but also involves challenges in neural control: transforming internal states ^14^ and external sensory cues into motor outputs that are **flexible**, **adaptive**, and **robust** in the face of uncertainty or conflicting drives^15–17^.

Each of these stages, from spatial encoding to motor execution, depends on specialized neural circuits. Much of our current understanding of spatial representations comes from studies in rodent models, where place cells, grid cells, and head-direction cells provide detailed maps of environmental and body-centered coordinates ^7,18–20^. However, the full navigation process also requires **decision-making and control systems** to convert these representations into movement. Investigating these later stages has been challenging in mammals due to the complexity and inaccessibility of distributed brain circuits.

In contrast, the fruit fly, Drosophila melanogaster, provides an opportunity to study navigation and motor control at the single-cell level. Although analogs of place and grid cells have not yet been identified in flies, the availability of powerful genetic tools and brain connectomes, such as FlyCircuit ^21,22^, hemibrain ^23^, and FlyWire ^24^, enables high-resolution circuit dissection. Recent studies have elucidated how flies encode head direction via a ring attractor mechanism ^25–30^, how synaptic plasticity contributes to flexible encoding ^31–33^, how wind, heading and motor direction are integrated through vector operation in the central complex ^34–37^, and how the central brain and the descending neurons generate motor responses ^38–43^.

Importantly, attention has begun to shift from representation to action. In addition to extensively studied visuomotor control of *Drosophila* ^44,45^, recent studies investigated goal-directed steering. Neurons known as PFL3 receive inputs encoding current heading direction (via the protocerebral bridge) and internal goal direction (via FC2 neurons) ^36^. By summing these signals, PFL3 neurons encode the angular difference between heading and goal, effectively computing a “directional error signal ^36^.” Their outputs then drive descending neurons (DNs), which in turn initiate turning movements to reduce this error and align the fly with its goal ^35,36^. This model provides a simple and elegant solution to the problem of steering, echoing the sensorimotor principles proposed by Valentino Braitenberg in his classic “Vehicles” ^46^.

However, real-world navigation involves far more than steering toward a static goal. Effective control systems must handle **conflicts, interruptions, and physical contingencies**. Aversive stimuli near the goal may require a temporary retreat. Obstacles may trigger detours. Injuries or body dynamics may distort motor responses. Sensory symmetry, such as when an aversive stimulus lies directly in front of the fly ^35^, may lead to neural stalemates. In such cases, the system must not only compute an error signal, but must **adapt, override, or resolve competing drives**. This requires a control architecture that is both goal-directed and **flexibly reconfigurable**.

In this study, we hypothesize that such flexible control emerges from circuit motifs within the lateral accessory lobe (LAL) ^47,48^, a brain region interposed between PFL3 and descending neurons. Using the Flywire and hemibrain connectome databases, we identify a network of direct, indirect, recurrent and feedforward pathways in the LAL. Through in-depth analysis and computer simulations, we find that these circuits implement two distinct steering modes: a **pro-goal response** that turns the fly toward a target, and an **anti-goal response** that turns it away from aversive stimuli near the goal. Furthermore, the circuit architecture can resolve symmetrical input conditions: rear stalemate and front stalemate, through mechanisms that amplify small perturbations into decisive actions.

By modeling these circuits, we demonstrate that LAL enables a compact yet powerful control strategy that balances goal pursuit with environmental contingencies. The architecture mirrors core principles of control theory and behavioral arbitration, such as comparator circuits, override relays, and layered behavioral hierarchies, and may represent a scalable strategy for robust action selection. These findings not only shed light on neural control in *Drosophila*, but also offer design principles for bio-inspired embodied agents with minimal yet adaptive neural systems.

## Results

### The goal-directed steering behavior and associated LAL neurons

We investigated the neural circuits underlying goal-directed steering behavior, in which a goal is initially offset from the fly’s head direction, requiring the fly to turn its body until it becomes aligned with the goal (Fig. 1A). Previous work identified PFL2 and PFL3 neurons as encoding the head-goal offset (*Δθ*) and projecting downstream descending neurons (DNs) that drive turning behavior (Fig. 1B).

**Fig. 1.**
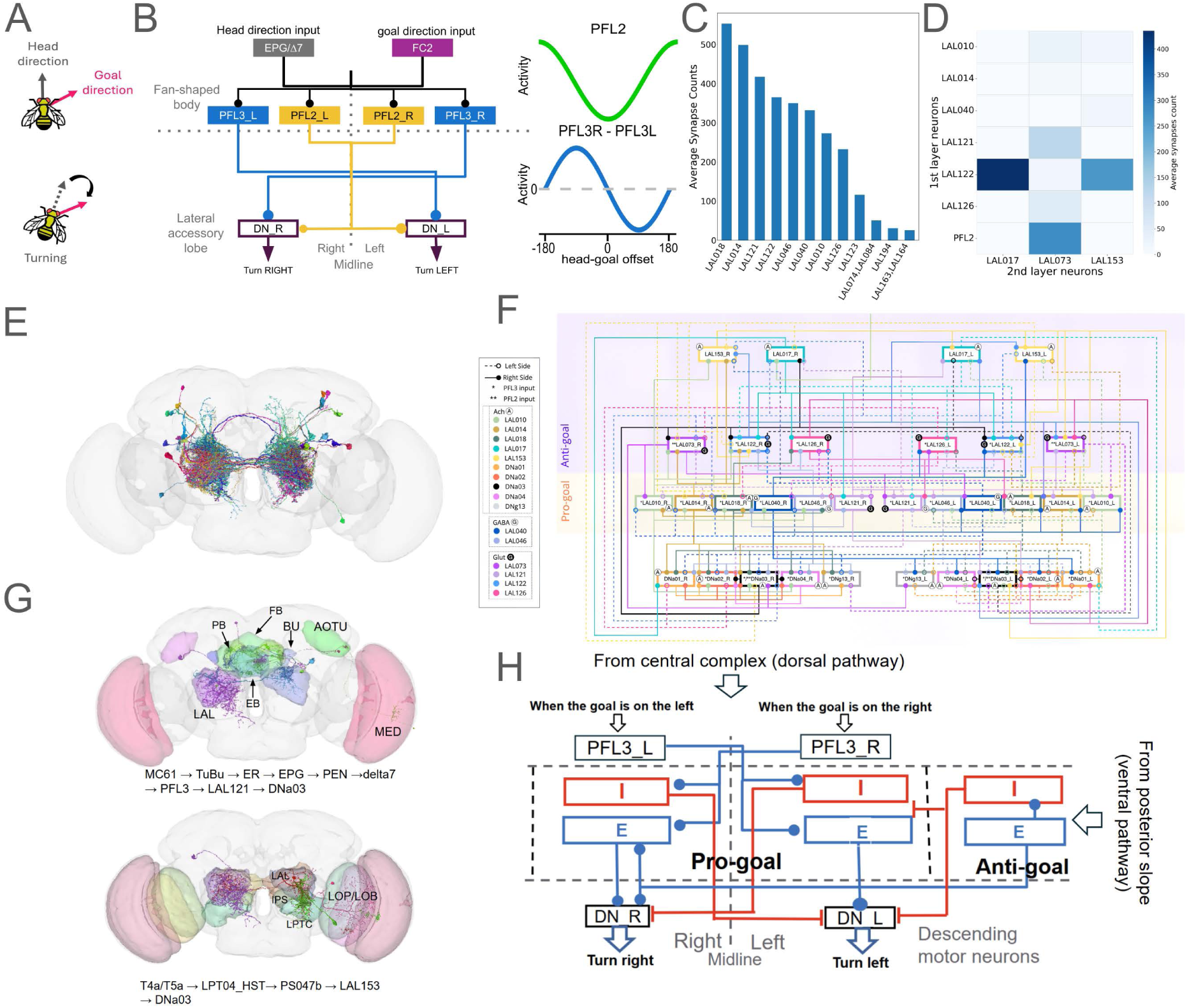
Goal-directed steering behavior and associated LAL control circuits in *Drosophila melanogaster*. **A.** Schematic of goal-directed steering behavior. A fruit fly rotates its body until its heading aligns with internally defined goal direction. **B.** (Left) Previously identified circuits underlying goal-directed steering ^35,36^. The circuits involve the fan-shaped body (FB) and lateral accessory lobes (LAL), with descending neurons (DNs) initiating body rotation. (Right) The input signals for heading and goal direction are encoded differently by PFL3 and PFL2 neurons, which represent the angular offset (Δ*θ*) between heading and goal. **C.** Average synapse counts between PFL3-connected LAL neurons (presynaptic) and DNa01-04 neurons (postsynaptic). Neurons with > 100 synapses were selected as the first set for modeling. **D.** A second set of LAL neurons that strongly interact with the first set. The color-coded matrix shows mean synapse counts between and within the two sets. **E.** Skeletons of neurons included in the model, reconstructed the Flywire database ^24^. **F.** Connectivity diagram of all modeled neurons based on Flywire databases. Each rectangle denotes a neuron type; connecting lines represent synaptic connections. **G.** Visual information reaches the LAL neurons through two main routes: a dorsal pathway (top) via the central complex and a ventral pathway (bottom) via the inferior posterior slope (IPS). The plots show representative signal paths from the medulla to descending neurons through each route. **H.** Based on their input pathways and connectivity, the LAL neurons can be grouped into two functional sub-circuits: the pro-goal circuit, which drives turning toward the goal, and the anti-goal circuit, which promotes turning away from it. Excitatory (E) and inhibitory (I) neuron groups are shown as simplified writing diagrams for visualization.

By analyzing connectome data, we found that the previously identified circuit provides only a partial picture of goal-directed steering control. Using the Flywire database, we identified a cluster of lateral accessory lobe (LAL) neurons that receive strong input from PFL2/3, and, in turn, form dense synaptic connections with the turning-related descending neurons DNa01-04. Among these, DNa02 is known to play a crucial role in mediating turning movements^35,38,40,49^, while DNa01, 03 and 04 were included in our analysis because of their strong connectivity with DNa02.

The selected LAL neurons, LAL018, 014, 121, 122, 046, 040, 010 and 126, constitute the first set of neurons in our proposed model (Fig. 1C). Cross-validation using the hemibrain database confirmed that most of these neurons are among the most strongly connected LAL neurons between PFL2/3 and DNa01-04, except for a few that project contralaterally (and Supplementary Fig. 1). Because the hemibrain dataset covers only one hemisphere of the brain (except for a few central complex neuropils), our model was primarily developed based on the bilateral connectome data from the Flywire.

We further identified additional LAL neurons that form strong synaptic connections with this first set. Although receiving no direct PFL2/3 input, the LAL017 and 153 form strong connections with LAL121, 122, 010 and 040. LAL073 receives inputs from PFL2 and connects with LAL121, 153 and 040. These three neurons (LAL017, 153 and 073) may therefore also participate in the steering control and were incorporated into the model (Fig. 1D).

The complete circuit modeled in this study, including input neurons (PFL2 and PFL3) and output descending neurons (DNs), consists of 12 PFL2 neurons, 24 PFL3 neurons, 22 LAL neurons (bilateral LAL 010, 014, 017, 018, 040, 046, 073, 121, 122, 126, 153), and 8 DNa neurons (bilateral DNa 01-04)(Fig. 1E). Note that LAL122 appears as a pair of neurons on each side. These neurons form 236 connections (Supplementary Table 1) when we consider only those with more than 5 synapses, to reduce potential errors in automated identification. The LAL neurons not only project feedforward to the descending neurons but also form extensive interconnects among themselves (Fig. 1F).

To further elucidate the functional organization of this circuit, we examined its upstream inputs. In addition to the known PFL3/PFL2 inputs, we found that several LAL neurons also receive inputs from the inferior protocerebrum superior IPS (Supplementary Fig. 2), which can be traced back to the optical lobe. Two distinct pathways were identified. The **dorsal pathway** originates in the medulla, and proceeds through AOTU, BU, EB, PB, and FB, before reaching the LAL (Fig. 1G top). The **ventral pathway** also starts from the medulla, but passes through LOP, LPTC, and IPS before arriving at the LAL (Fig. 1G bottom). The terms “dorsal” and “ventral” refer to their relative anatomical positions rather than implying correspondence to the dorsal/ventral visual pathways described in mammalian cortices.

These two pathways target largely distinct groups of LAL neurons, suggesting a division into two functional subcircuits. Through detailed connectivity analysis and computer simulations, we found that the full network can be separated into two modules: the **pro-goal circuit,** primarily receiving inputs from the dorsal pathway, and the **anti-go circuit**, primarily receiving inputs from the ventral pathways (Fig. 1H). Excitatory neurons in the pro-goal circuit facilitate turning toward the goal by exciting ipsilateral excitatory or descending neurons, while inhibitory neurons suppress contralateral excitatory or descending neurons. In contrast, the anti-goal circuit exhibits an opposite connectivity pattern, suppressing the pro-goal response and promoting turning away from the goal direction (Fig. 1H).

For clarity, we renamed each neuron according to its upstream input pathway (V for ventral; D for dorsal) and the predicted functional polarity (E for excitatory and I for inhibitory), as annotated in the Flywire database. The correspondence between Flywire neuron ID and the functional names used in this paper is summarized in Table 1.

**Table 1:** Neuron ID (Flywire) and Functional Name (this paper) Mapping.

| Neuron ID in FlyWire | ID in the present paper | Circuit classification / Tentative function | Neuron ID in FlyWire | ID in the present paper | Circuit classification / Tentative function |
| --- | --- | --- | --- | --- | --- |
| LAL010 | DE1 | pro-goal / excitation | LAL073 | DI4 | pro-goal / inhibition |
| LAL014 | DE2 | pro-goal / excitation | LAL153 | VE1 | anti-goal / excitation |
| LAL018 | DE3 | pro-goal / excitation | LA017 | VE2 | anti-goal / excitation |
| LAL040 | DI1 | pro-goal / inhibition | LAL122 | VI1 | anti-goal / inhibition |
| LAL121 | DI2 | pro-goal / inhibition | LAL046 | VI2 | anti-goal / inhibition |
| LAL126 | DI3 | pro-goal / inhibition |  |  |  |

### The pro-goal circuit

We first analyzed the subcircuit that receives PFL2/3 inputs and promotes turning toward the goal direction, which we refer to as the pro-goal circuit. It includes DE1 (LAL010), DE2 (LAL014), DE3 (LAL018), DI1 (LAL040) DI2 (LAL121), DI3 (LAL126) and DI4 (LAL073) (Fig. 2A). DE1-3 are tentatively cholinergic neurons that receive the contralateral PFL3 inputs and project to the ipsilateral descending neurons (DNs). DE1 and DE2 form weak reciprocal connections and both project to DE3. These three neurons form an **excitatory core (EC)** of the pro-goal circuit. In contrast, DI1 is likely GABAergic, and DI2-4 are tentatively glutamatergic inhibitory neurons. They also receive contralateral PFL3 inputs but inhibit contralateral DNs and EC. This connectivity suggests a **push-pull mechanism:** when one hemisphere receives strong PFL3 input, the excitatory core amplifies the excitation on the ipsilateral DNs while the inhibitory neurons suppress the contralateral side.

**Fig. 2.**
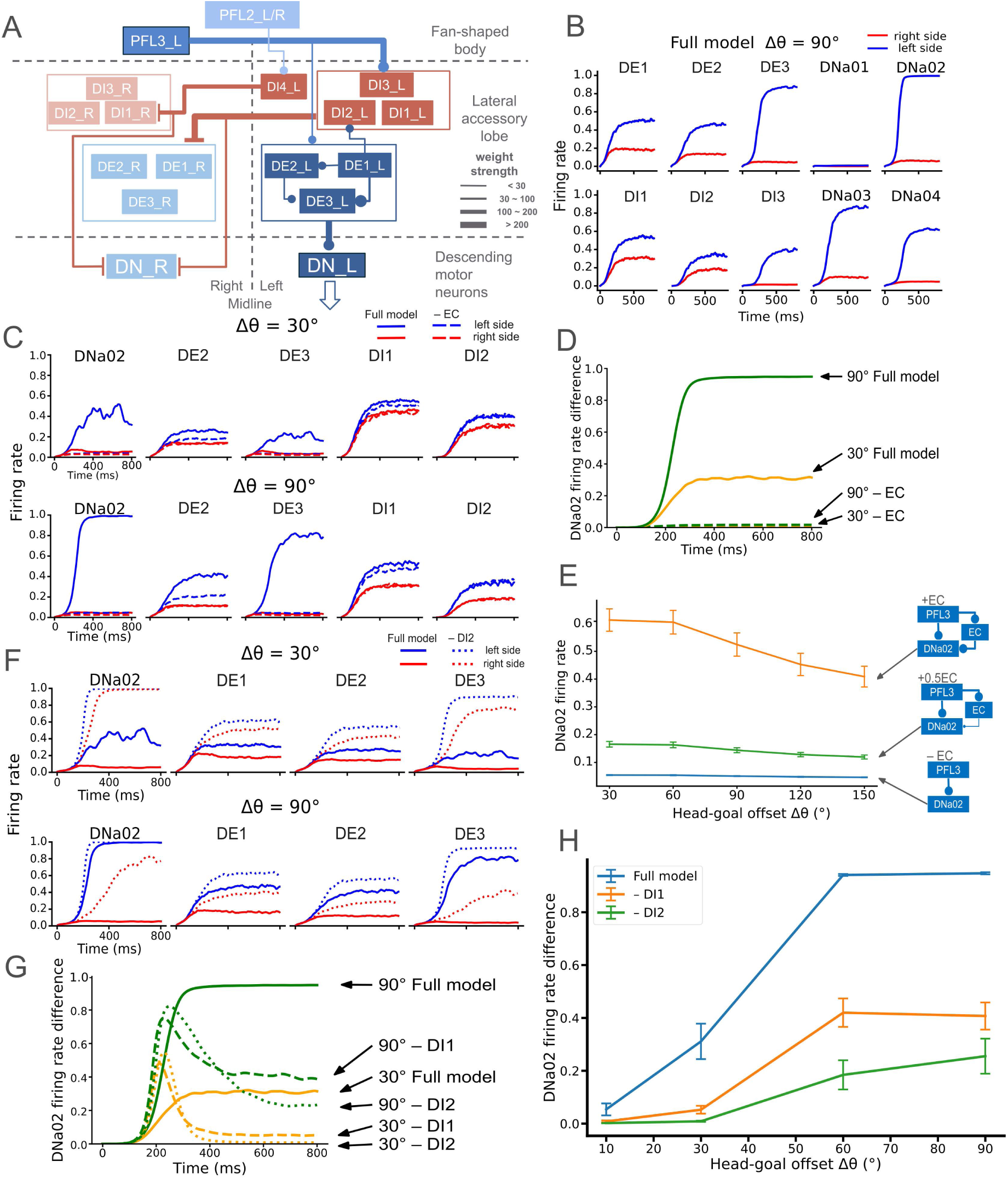
The pro-goal circuit. **A.** Schematic of neurons and major connections in the pro-goal circuit. DE1 (LAL010), DE2 (014) and DE3 (018) form an excitatory core (EC) that drives the ipsilateral descending neurons, whereas DI1 (LAL040) and DI2 (121) provide contralateral inhibition to the opposite pro-goal circuit and descending neurons. The circuit is bilaterally symmetric; only the left side is shown for clarity. **B.** Mean neuronal responses at *Δθ*=90° (goal to the left) averaged over 50 trials. The descending neuron DNa02 exhibits higher firing rates on the left than on the right, indicating a left-turning command. **C.** Mean responses of the full model (solid curves) and the model without the excitatory core (-EC; dashed curves) at *Δθ*=30° (top) or 90° (bottom) averaged over 50 trials. Removing the EC markedly reduces the left-right firing-rate difference across multiple neurons, including DNa02. **D.** Mean firing-rate difference between left and right DNa02 neurons (*Δr*_*DNa*02_) at *Δθ*=30° and 90° for the full model (solid curves) and no EC condition (-EC; dashed curves), averaged over 50 trials. **E.** Mean *Δr*_*DNa*02_ (50 trials) as a function of *Δθ* in an isolated unilateral circuit containing only PFL3, EC and DNa02, tested with full (+EC), half (+0.5EC) or no 0% (-EC) EC output strengths. EC activity enhances DNa0 responses across all *Δθ* values. **F.** Mean neuronal responses (50 trials) in the full model (solid) and without DI2 (-DI2; dashed). Most neurons show higher overall firing but reduced left-right asymmetry with DI2 is removed. **G.** Mean Δ*r*_*DNa*02_ in the full model and after removal of DI1 or DI2. **H.** As in E, but for the full model and DI1 or DI2 removed. Eliminating either inhibitory neuron type significantly reduces *Δr*_*DNa*02_ across all *Δθ* values.

To explore the circuit’s dynamics, we built a computational model using the identified connectivity under an open-loop configuration, where inputs (goal-head offset) were fixed and not influenced by DN outputs. The open-loop setup allowed us to examine the steady-state responses and clarify each neuron’s contribution. When PFL3 fire rates represented a 90° goal-head offset (*Δθ*=90°, goal on the left), the circuit rapidly generated lateralized activity: within ∼100 ms, DNa02_L became more active than DNa02_R, initiating a left-turning response that would reduce the offset (Fig. 2B and Supplementary Fig. 3).

We next examined the functional role of the excitatory core (DE1-DE3). Because PFL3 neurons already innervate DNs directly, we asked why an additional excitatory relay exists. Removing the EC drastically reduced the firing-rate difference between left and right DNa02 (Δ*r*_*DNa*02_) (Fig. 2C and Supplementary Fig. 4), indicating that the EC provides essential amplification. This effect was robust across a wide range of Δ*θ* values (Fig. 2D). Note that the relative connection strengths were determined by the number of synapses from the database; therefore, the importance of EC did not result from the choice of tuning parameters but was the consequence of observed connectivity. To isolate its contribution, we simulated a minimal unilateral circuit containing only the EC, its upstream PFL3, and its downstream DNa02. In this simplified model, direct PFL2/3 input alone was insufficient to activate DNa02, whereas adding the EC reinstated robust activation (Fig. 2E). Thus, the excitatory core acts as an auxiliary drive that strengthens the pro-goal output.

The circuit also contains contralaterally projecting inhibitory neurons (DI1-DI4). Because turning speed depends on the firing rate difference between the left and right DNa02 neurons (Δ*r*_*DNa*02_), excitation on one side must be complemented by suppression on the other.

Removing DI neurons abolished this push-pull effect: both DNa02_L and DNa02_R became highly active, and Δ*r*_*DNa*02_ dropped significantly (Fig. 2F and Supplementary Fig. 5). We performed 50 trials and verified that *Δr*_*DNa*02_ was significantly reduced without DI1 or DI2 (Fig. 2G & 2H). Our tests showed that removing DI3 or DI4 produced similar, but much smaller, effects on DN02 responses (Supplementary Fig. 6). We therefore focused subsequent analyses on DI1 and DI2. Together, these results show that the pro-goal circuit amplifies asymmetries in PFL3 input through a coordinated “push-pull” architecture, in which the excitatory core drives ipsilateral DNs and inhibitory neurons suppress contralateral activity to promote turning toward the goal.

### Rear Stalemate

The pro-goal circuit exhibits an inherent limitation: turning becomes sluggish when the goal lies behind it (*Δθ* ∼180°), and completely ceases when the goal is exactly opposite to the head direction (*Δθ* =180°). We refer to this condition as the “Rear Stalemate” (Fig. 3A). It arises because the firing rate difference between PFL3_R and PFL3_L declines when *Δθ* >90° and reaches zero at *Δθ* =180° (Fig. 1B). The rear stalemate was investigated recently and another neuron group, the PFL2s, was shown to contribute to resolving this problem^35^. Unlike PFL3 neurons, which project contralaterally, PFL2_R and PFL2_L project bilaterally to several LAL neurons and DNa03, with firing rates that peak at *Δθ* =180° and decrease monotonically as |*Δθ*| decreases (Fig. 1B). This bilateral input provides an extra drive that compensates for weak PFL3 activity and accelerates the turning rate at *Δθ* ∼180°.

**Fig. 3.**
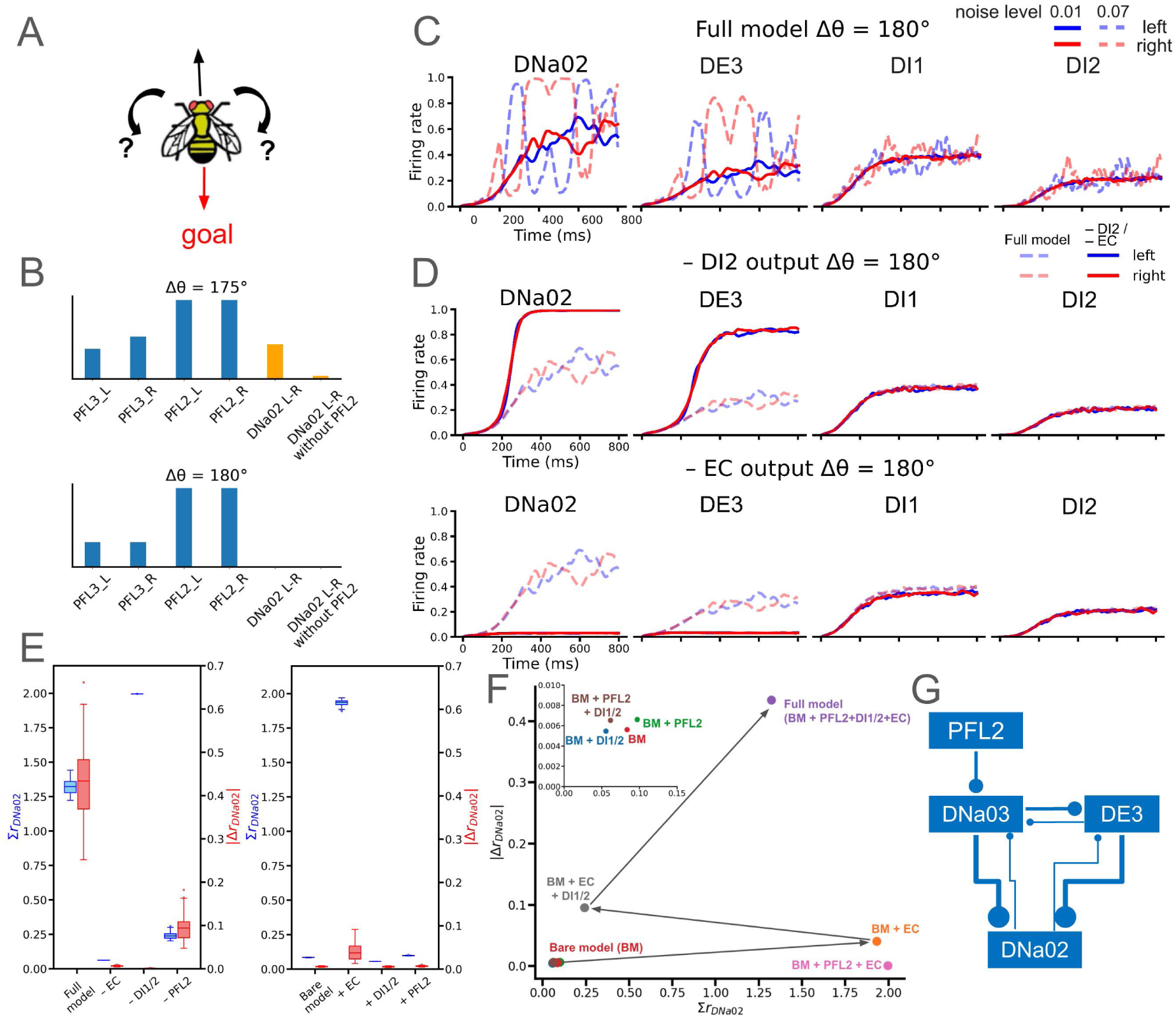
Rear Stalemate and how the circuit resolves it. **A.** Schematic illustrating Rear Stalemate, which occurs when the goal lies directly behind the head (*Δθ*=180°). At this offset, PFL3 inputs and DNa neuron responses are symmetric, producing no turning. **B.** Schematics illustrating the effect of PFL2 on Rear Stalemate. Top, when *Δθ* = 175°, PFL2 amplifies small asymmetries in PFL3 firing, increasing *Δr*_*DNa*02_ and promoting turning. Bottom When *Δθ*=180°, both PFL2 and PFL3 inputs are perfectly symmetric, so PFL2 exerts little effect on Δ*r*_*DNa*02_., **C.** Example trials showing the impact of noise at *Δθ*=180°. With weak Gaussian noise (s.d.= 0.01, solid curves), the system occasionally develops large |*Δr*_*DNa*02_|, indicating successful escape from the stalemate. Strong noise (s.d. = 0.07, dashed) enhances this effect and accelerates symmetry breaking. **D**. Representative trials demonstrating the the essential roles of the excitatory core (EC, top) and inhibitory neuron DI2 (bottom). Removing either component nearly abolishes |*Δr*_*DNa*02_|. **E.** Left, effects of removing PFL2, EC, or DI1/2 from the full model (averaged across 50 trials). The intact circuit yields the largest |*Δr*_*DNa*02_|, whereas deleting any single component diminishes symmetry breaking. Right, results from a “bare model” lacking all three components, with each re-added individually. **F**. Summary plot of *Δr*_*DNa*02_ vs. *∑r*_*DNa*02_. Arrows indicate how sequentially adding EC, DI and PFL2 to the bare model (BM) maximizes *Δr*_*DNa*02_ (see text for details). **G.** Connectivity between PFL2 and DE2 (part of EC) subcircuits. Their interaction produces a nonlinear amplification that drives stronger excitation of DNa02 than either pathway alone.

However, even with PFL2 input, the stalemate is not fully resolved, because at *Δθ*=180°, both PFL2 and PFL3 inputs are perfectly symmetric (Fig. 3B). To break this symmetry, DNa02 and DNa03 must receive slightly unbalanced inputs. As expected, adding noise introduces such asymmetry: higher noise levels produced larger |*Δr*_*DNa*02_|, effectively restore motion (Fig. 3C and Supplementary Fig. 7). Yet noise alone was insufficient. We found that additional circuit components, including EC and DIs, were crucial for robust symmetry breaking. Removing EC or DI2 markedly reduced |*Δr*_*DNa*02_| (Fig. 3D and Supplementary Fig. 8).

We systematically tested the contributions of PFL2, EC and DI to resolving Rear Stalemate by measuring the mean sum, *∑r*_*DNa*02_ (= *r*^*R*^_*DNa*02_ + *r*^*L*^_*DNa*02_), and difference |*Δr*_*DNa*02_| (= |*r*^*L*^_*DNa*02_ − *r*^*R*^_*DNa*02_|) across 50 trials under elavated noise (s.t.d. = 0.07)(Fig. 3E). Loss-of-function tests showed that removing any of the three components (EC, DI2, PFL2) decreased *Δr*_*DNa*02_, indicating reduced ability to escape the stalemate (Fig. 3E left). Conversely, gain-of-function tests, adding each component to a bare mode (BM) lacking EC, DI and PFL2, revealed that none of them alone could reproduce the large |*Δr*_*DNa*02_| of the full model (Fig. 3E right). Together, these analyses show that symmetry breaking requires the combined action of PFL2, EC and DI.

We visualized this cooperative effect by sequentially adding components to the bare model (Fig. 3F).

1. Adding EC strongly increased total DNa02 activity (*∑r*_*DNa*02_) but produced little ^|*Δr*^*DNa*02^|.^
2. Adding DI (BM+EC+DI) slightly increased |*Δr*_*DNa*02_| but reduced *∑r*_*DNa*02_ due to inhibition from DI, limiting the maximum possible difference because it is always smaller than |*Δr*_*DNa*02_|.
3. Added PFL2 (full model) restored *∑r*_*DNa*02_ and allowed DI to further enhance |*Δr*_*DNa*02_|.

Two excitatory sources, EC and FPL2, were both essential. When only one excitatory group was present (BM+PFL2+DI or BM+EC+DI), *∑r*_*DNa*02_ became too small to sustain a strong difference. Their co-existence produced a “nonlinear enhancement”: |*Δr*_*DNa*02_| with both groups was far greater than the sum of their individual effects. This nonlinearity emerged from circuit interactions: PFL2 projected to DNa03, which excited DE2 (part of EC); DE2 then projected to DNa02, the primary driver of turning (Fig 3G). Thus, PFL2 activation potentiated EC output, leading to strong DNa02 activation. Without EC, PFL2’s effect on DNa02 was greatly diminished.

In summary, Rear Stalemate represents a symmetry-induced indecision and halts steering. It is overcome by a combination of noise-driven symmetry breaking and the cooperative action of three components, PFL2, EC and DI, that together accerlerate winner selection and reinitiate turning.

### The anti-goal circuit

We identified another subcircuit that projects to descending neurons (DNs) in a manner opposite to the pro-goal circuit, which we term the “anti-goal circuit”. It comprises VE1 (LAL153, cholinergic), VE2 (LAL017, cholinergic), VI1 (LAL122, glutamatergic), and VI2 (LAL046, GABAergic), which receive little or no inputs from PFL2/3 (Fig. 4A). Instead, VE1, VE2 and VI2 receive strong inputs unilateral input from the inferior posterior slope (IPS), likely carrying visual information^38,50^. VE1 projects to the contralateral DN (Fig. 4A, path 1), so a visual signal on one side may evoke a turn away from it. VE1 also projects to contralateral VE2 and VI1 (Fig 4A, path 2), and VI1 in turn inhibits DNs ipsilateral to the activated VE1 (Fig. 4A, path 3). VI2 receives ipsilateral IPS input (similar to VE1) and inhibits ipsilateral DNs (Fig 4A, path 4). Together, these interactions amplfy VE1-driven turning.

**Fig. 4.**
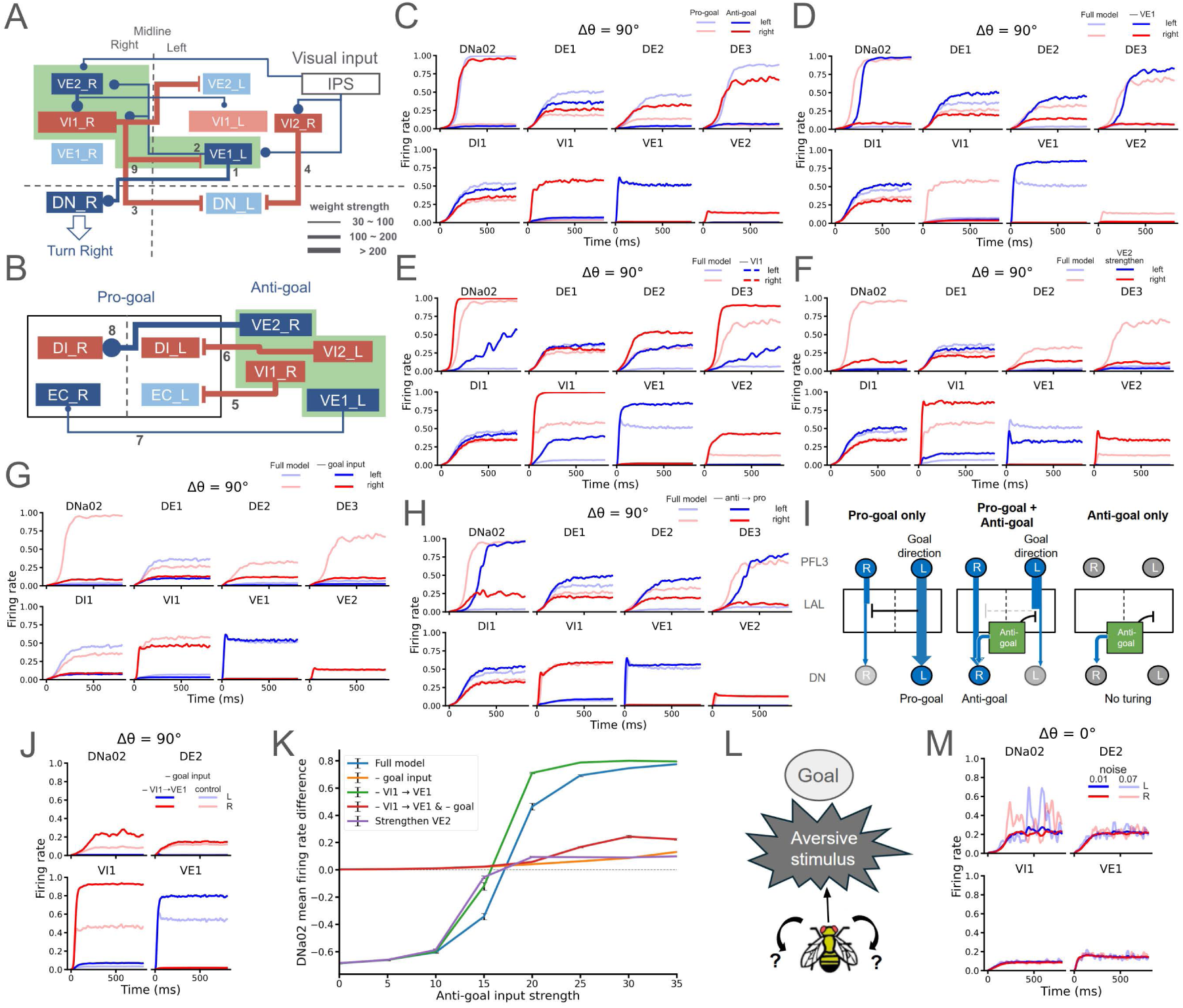
The anti-goal circuit and its functional mechanisms. **A.** Schematics of neurons and major connections in the anti-goal circuit, composed of tightly coupled VE1 (LAL153), VE2 (LAL017), VI1 (LAL122) and VI2 (LAL046) neurons. When activated by unilateral input from inferior posterior slope (IPS), VE1 excites contralateral descending neurons (DNs), whereas VI1 and VI2 inhibit DNs on the ipsilateral side of the IPS input. **B.** Major projections from the anti-goal to the pro-goal neurons. These pathways suppress ipsilateral pro-goal activity and thereby facilitate contralateral (anti-goal) turning. Numbers in **A** & **B** label the important pathway explained in the text. **C.** Neural activity during anti-goal turning (pro-goal *Δθ* =90°, plus anti-goal input) compared with pro-goal turning alone. **D-F.** Same as **C** but with VE1 removed, VI1 removed or VE2’s input strengthened, respectively. Each manipulation demonstrated the specific contribution of these neurons to the anti-goal response. **G.** Neural activity during the anti-goal task when pro-goal input is absent. Without pro-goal drive, the anti-goal circuit alone fails to elicit turning. **H.** Same as **G** but with anti-goal-to-pro-goal connections removed. Eliminating these cross-links also abolishes anti-goal turning. **I.** Schematic of the conditioned anti-goal mechanism. Left, with only pro-goal input, the active (left) pro-goal neurons inhibit the contralateral (right) side. Middle, when anti-goal input is applied, the anti-goal circuit suppresses the left pro-goal response, releasing the right side and enhancing a rightward turn. Right, without pro-goal input, the anti-goal circuit alone cannot initiate turning. **J.** The VI1→VE1 feedback connection is critical for this conditional behavior. Removing it partially rescues anti-goal responses in the absence of pro-goal input. **K.** Systematic tests of anti-goal responses under various conditions: removing pro-goal input, removing VI1→VE1 connections, removing both, or enhancing VE2. The model failed to generate anti-goal response without the pro-goal input (orange curve) but the response partially recovered when the VI1→VE1 feedback was removed. **L.** Front Stalemate: when both the aversive stimulus and goal are directly in front of the fly, bilateral anti-goal circuits receive identical inputs, producing a stalemate. **M.** The anti-goal response can still emerge in this stalemate condition if the noise level is sufficiently high.

The anti-goal also interacts with the pro-goal circuit. VI1 and VI2 inhibit EC and DI neurons contralateral to the anti-goal direction (Fig 4B, paths 5 & 6), whereas VE1 and VE2 activate EC and DI on the ipsilateral side (Fig 4B, paths 7, 8). These cross-connections allow the anti-goal circuit to override the pro-goal circuit when the two conflict. Additionally, VI1 provides feedback inhibition contralateral VE1, likely to prevent it from overactivation in an anti-goal response (Fig. 4A, path 9).

We tested these predictions in simulation by presenting a left IPS input, simulated by activating left VE1, VE2 and VI2, while also presenting a goal on the left (*Δθ*=90°) of the head direction. The unilateral activation from IPS generated higher right-side DNa02 activity, representing an anti-goal response (Fig. 4C and Supplementary Fig. 9). Lesioning VE1 (removing all its outputs) abolished this reversal (Fig. 4D and Supplementary Fig. 10) confirming its key role. Removing VI1 weakened the effect, producing only a small Δ*r*_*DNa*02_ (Fig. 4E and Supplementary Fig. 11). Manipulating VE2 revealed that it primarily acts as a brake: overactivating it reduced *Δr*_*DNa*02_, whereas lesioning it had little effect*Δr*_*DNa*02_(Fig. 4F and Supplementary Fig. 12).

Unexpectedly, the anti-goal response required concurrent pro-goal input from PFL2 and PFL3. When these inputs were removed, identical IPS activation failed to elicit an anti-goal response (Fig. 4G Supplementary Fig. 13). Intuitively, an anti-goal input needs to overcome the pro-goal response, thus its removal should facilitate the reversed turn. To explore this paradox, we disconnected the anti-goal-to-pro-goal links so that the two circuits operated independently (Fig. 4H and Supplementary Fig. 14). The anti-goal circuit alone could not outcompete the pro-goal circuit, showing that the cross-connections between the two circuits are essential. This counterintuitive phenomenon stems from an intricate mechanism in which the anti-goal circuit recruits the pro-goal circuit and reverses it (Fig. 4B and Fig. 4I). During normal pro-goal turning, e.g., leftward, stronger left-side excitation suppresses the right side via DI-mediated inhibition (Fig. 4I, left). When anti-goal activation is required, the anti-goal neurons inhibit the left pro-goal side, releasing inhibition on the right and adding a small excitation to right-side DNs, thus flipping the steering direction (Fig. 4I, middle). Without pro-goal input, this small excitation alone cannot drive turning (Fig. 4I, right).

One may expect that sufficiently strong anti-goal input could act alone, but the VI1-->VE1 feedback prevents such runaway activation (Fig. 4A, path 9). Removing this connection in the no-pro-goal condition partially restored the anti-goal response (Fig. 4J and Supplementary Fig. 15). Systematic tests varying anti-goal input strength confirmed these roles (Fig. 4K): removing VI1-->VE1 lowered the threshold for triggering an anti-goal response, whereas removing pro-goal input abolished DN activation altogether. Combining both manipulations partially rescued the response, indicating that the VI1 feedback imposes conditional gating. Overactivating VE2 eliminated anti-goal responses at all input levels, consistent with its role as a brake.

In summary, the anti-goal circuit operates through three coordinated mechanisms:

1. **Direct excitation and inhibition** of DNs to bias turning.
2. **Recruitment and reversal** of the pro-goal circuit. Specifically, the anti-goal circuit hijacks the pro-goal circuit by repurposing the pro-goal circuit and generating strong output in the opposite direction.
3. **Brake / output limiter**. The anti-goal excitatory neurons recruit inhibitory feedback, creating a self-limiting loop.

Functionally, the anti-goal circuit behaves like a “low-power override controller” in engineering systems: it does not directly command movement but instead reconfigures or inverts the main controller. When the pro-goal circuit is inactive (i.e., no goal is presented), there is nothing to override.

### Front Stalemate

Similar to the pro-goal circuit, the anti-goal circuit also encounters a symmetry problem: the front stalemate. When both the goal and the aversive stimulus are perfectly aligned with the head direction, the two sides of the anti-goal circuit receive inputs of equal strength, preventing directional bias (Fig. 4L). We tested this condition in simulations and found that, despite the symmetric input, the circuit could still overcome the pro-goal attraction and generate an anti-goal response when the noise level was sufficiently high (Fig. 4M and Supplementary Fig. 16).

### The closed-loop simulations

After analyzing the circuit dynamics in open-loop conditions, we next asked whether the LAL circuit can perform the proposed pro-goal and anti-goal functions in a closed-loop setup, where the circuit output (DN activity) updates the fly’s heading, which in turn modifies the PFL2/3 inputs (Fig. 5A; see Materials and Methods).

**Fig. 5.**
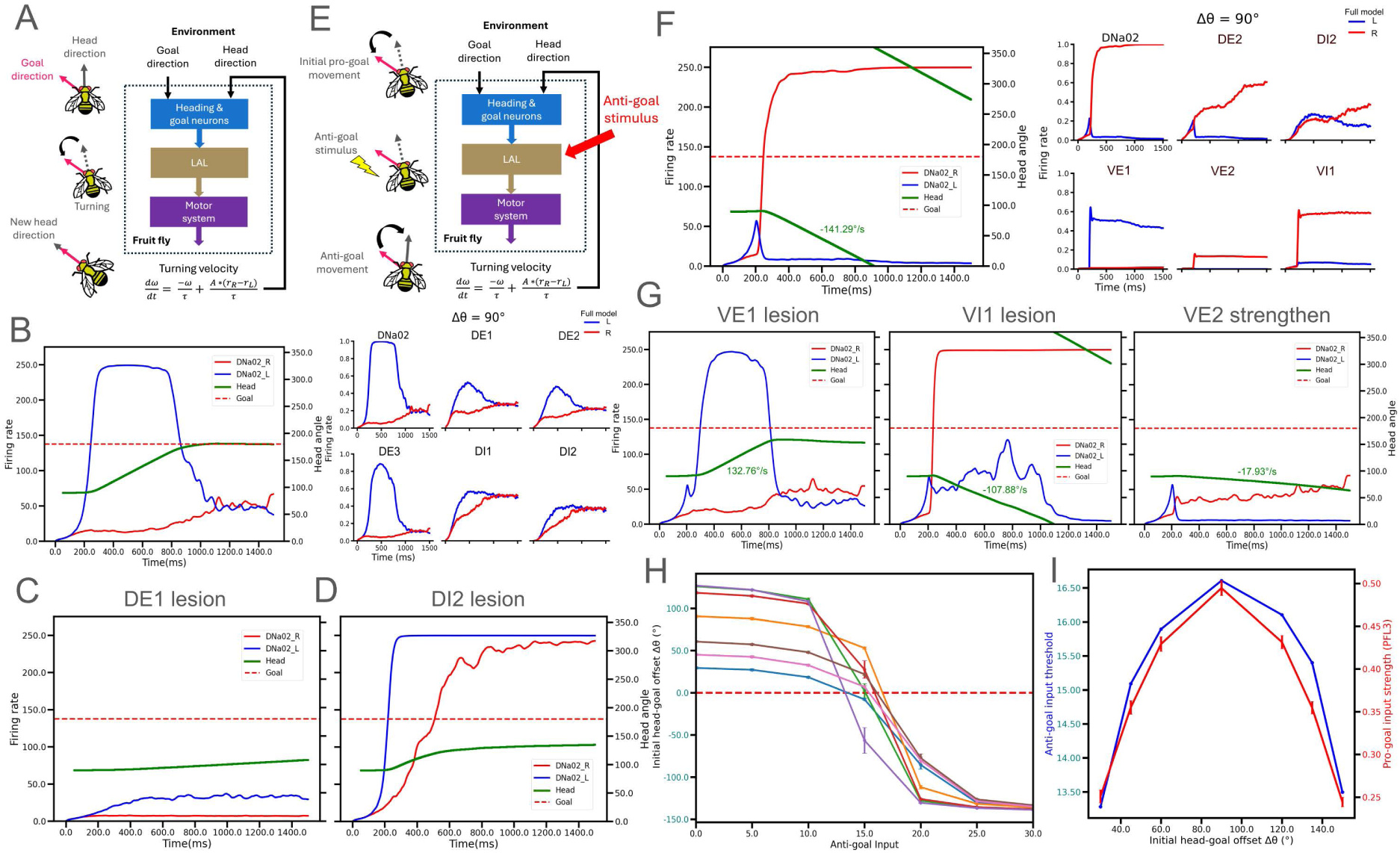
Closed-loop simulations of pro-goal and anti-goal behaviors. **A.** Block diagram of the closed-loop pro-goal simulations. At each time step, model output (DNa02 activity) is converted to angular velocity, integrated to update the fly’s heading, fed back as input for the next time step. **B.** Left. example closed-loop simulations of pro-goal turning (initial *Δθ* = 90°). The difference between the left (blue) and right (red) DNa02 activity drives body rotation until the heading (green) aligns with the goal (dashed line). Right, riring-rate traces of representative neurons in the pro-goal circuit. **C & D.** Example simulations with DE1 and EI2 lesioned, respectively. Both lesions markedly reduced pro-goal turning speed. **E.** Block diagram of the closed-loop anti-goal simulations. **F.** Left, example anti-goal simulations. After 200ms of pro-goal drive (initial *Δθ* = 90°), unilateral IPS input triggered a rapid turn away from the goal (turning rate shown in green). Right, firing-rate traces of key neurons in the anti-goal circuit. **G.** Effects of key neurons in anti-goal tests. Left, lesioning VE1 abolished the anti-goal response, producing pro-goal turning instead. Middle, lesioning VI1 did not abolish but moderately slowed the reversal. Right, strengthening VE2 output greatly reduced the anti-goal response. **H.** Model responses for various initial *Δθ* (color-coded) under different anti-goal input strengths. Anti-goal turning was triggered only when input exceeded a threshold. **I.** Threshold anti-goal input values depended on the initial *Δθ*, closely following the strength of the corresponding pro-goal drive from PFL3.

We first tested the system in a pro-goal task with *Δθ* = 90°. The circuit successfully aligned the heading with the goal within ∼1s, consistent with the behavioral timescale of real flies (Fig. 5B, Supplementary Fig. 17). The initial large PFL3 input generated strong *Δr*_*DNa*02_, which decreased as the fly approached the goal and reached zero when *Δθ* = 0. To assess the contribution of each component in this closed-loop context, we performed lesion tests. Removing any excitatory core (EC) neuron caused slower rotation and undershooting, while removing lateral-inhibition neurons (DI1 and DI2) produced similar behavioral effects (Fig. 5C,D; Supplementary Fig. 18). However, the underlying mechanisms differed: without EC, both DNa02 neurons were weakly activated, leading to small Δ*r*_*DNa*02_; without inhibitory neurons, both DNa02s remained active, again reducing *Δr*_*DNa*02_ through failed suppression.

We then performed the anti-goal task (Fig. 5E). Each trial began with a pro-goal turn (*Δθ* = 90°). After 200ms, unilateral IPS input was applied, which reversed the turning direction as predicted (Fig. 5F, Supplementary Fig. 19). Lesioning VE1 completely abolished the reversal, confirming its key role (Fig. 5G left; Supplementary Fig. 19A). Removing VI1 moderately reduced rotational speed (-107.88 °/s vs. -141.29 °/s in the control), while overactivating VE2 by tripling its IPS input drastically slowed turning (Fig. 5G middle & right; Supplementary Fig. 20B, C). These results agree with the open-loop tests (Fig. 4D-F).

To probe the interaction between goal and anti-goal drives, we varied the initial goal-head offset (*Δθ*) and IPS input strength (Fig. 5H). The model could trigger an anti-goal rotation at any Δ*θ*, but only when the IPS input exceeded a threshold. This threshold depended on the initial Δ*θ*, scaling with the strength of the pro-goal drive, which is approximatedly proportional to |*r*_*PFL*3*R*_ − *r*_*PFL*3*L*_ | (Fig. 5I). Thus, anti-goal activation requires a counter-signal strong enough to overcome the existing pro-goal momentum, a form of conflict resolution between competing control modules.

We next tested Rear and Front Stalemate under the closed-loop conditions. Starting from *Δθ* = 180° in the pro-goal task, the full model successfully broke the symmetry and initiated a pro-goal rotation within a few seconds (Fig. 6A; Supplementary Fig. 21). Lesioning EC, DI1/DI2 or PFL2, individually or in pairs, significantly delayed the reaction time, defined as the lantency to a 1.5° heading change after input onset. Removing one or any two of the components led to a significant increase of the reaction time (Fig. 6B), consistent with open-loop findings. For the front stalemate, when both goal and aversive stimuli were aligned with the head, the model initiated a turn away from the stimulus within a few hundred milliseconds (Fig. 6C, Supplementary Fig. 22). Lesion effects again matched the open-loop predictions Fig. 6D).

**Fig. 6:**
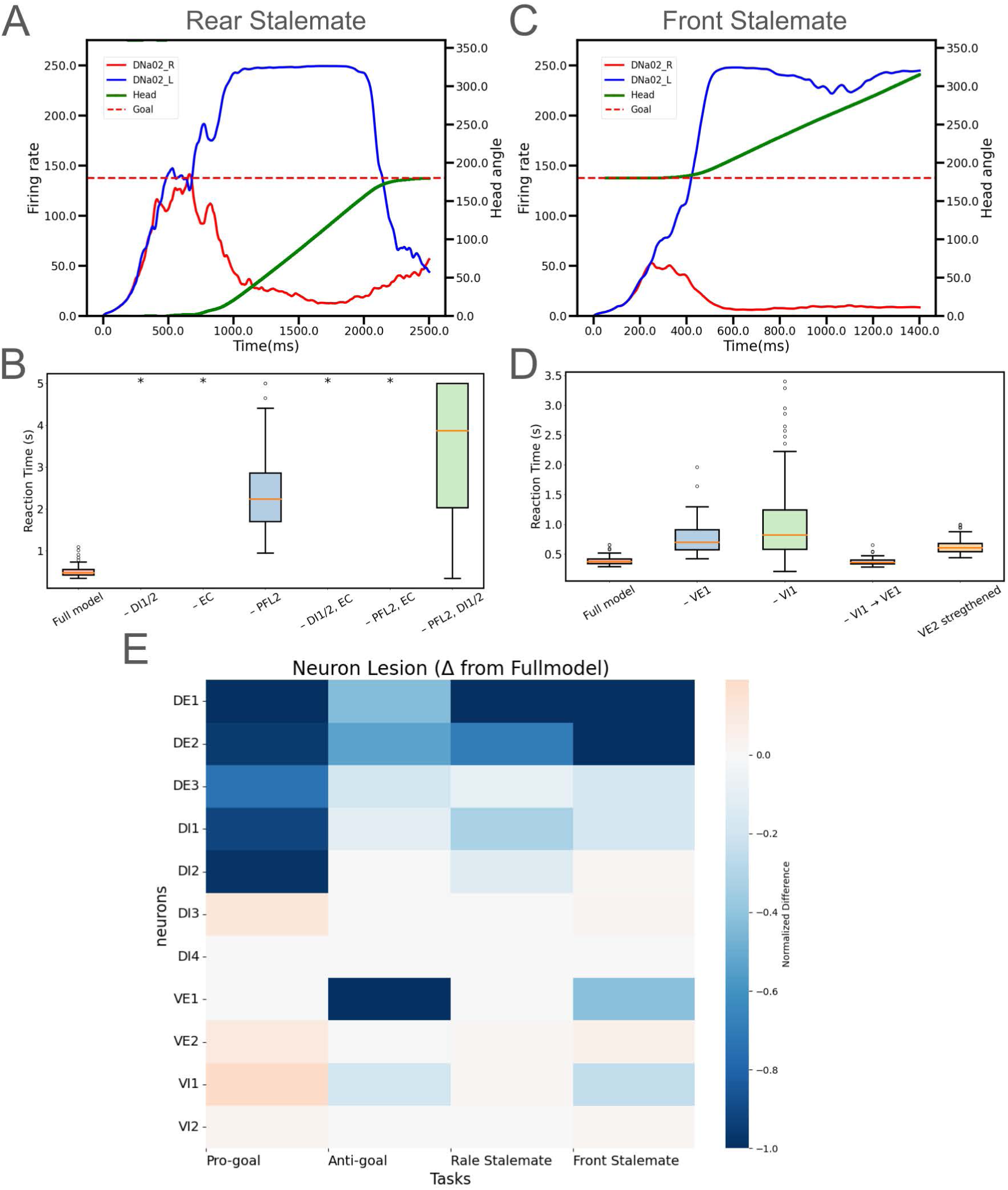
Closed-loop simulations of Rare & Front Stalemate conditions. **A.** Example closed-loop simulation of Rear Stalemate. The model resolved the stalemate within ∼500 ms and then rapidly turned toward the goal. **B.** Lesion tests of the key pro-goal components in Front Stalemate (100 trials per condition). Removing EC, DI or PFL2, or any two of them led to a greatly prolonged pro-goal reaction time. **C.** Example closed-loop simulation of Front Stalemate. The model overcame the symmetric input within ∼400 ms and turned away from the goal direction. **D.** Lesion tests of the key anti-goal components in Rear Stalemate (100 trials per. Removing VE1, VI1, or strengthening VE2 output connections increased the anti-goal reaction time. Removing the VI1 --> VE1 feedback, however, produced little change, consistent with Fig 4J & 4K. **E.** Systematic lesion analysis of all neuron types across pro-goal, anti-goal, Rear Stalemate, and Front Stalemate tasks. For each condition (30 trials, open-loop), we computed the difference in mean Δ*r*_*DNa*02_ between lesion and control (full) models. Results are color-coded: Positive values indicate enhanced responses, and negative values indicate reduced performance.

Finally, we conducted a systematic lesion analysis across all neuron types and tasks: pro-goal, anti-goal, rear stalemate and front stalemate, and measured the change in Δ*r*_DNa02_ relative to the intact model (Figure 6E). The results confirmed our functional classification. Notably, removing the EC (DE1-DE3) weakened responses in all tasks, including the anti-goal and front stalemate. This finding supports our earlier conclusion (Fig 4 G-I): the anti-goal circuit relies on recruiting an active pro-goal network to invert its output, and without EC activation, this recruitment, and hence anti-goal turning, cannot occur efficiently.

## Discussions

The lateral accessory lobe (LAL) of the *Drosophila* brain has been considered a hub for transforming sensory and goal-related input into descending motor commands. By analyzing the connectome and functional motifs within the LAL, we uncovered a set of circuit mechanisms that underlie flexible, context-dependent motor control, including mechanisms for pursuing internal goals and avoiding external threats. Our findings reveal two distinct yet interrelated neural pathways: the pro-goal circuit, which drives goal-directed steering, and the anti-goal circuit, which turns the body away from the goal. We also identified two challenge conditions: the Rear Stalemate and Front Stalemate, in which the system must resolve symmetry-induced indecision to produce a motor response. These motifs provide a computationally efficient, dynamically adaptive framework for action selection, and resonate with well-established architectures in control theory, digital systems, robotics and even molecular biology.

### Pro-goal circuit: Amplification without persistence

The pro-goal circuit receives visual input via the dorsal pathway and operates as a transient amplifier of directional intent. The bilateral set of pro-goal neurons excites ipsilateral descending neurons (DNs) and inhibits their contralateral counterparts, forming a push-full loop. This motif is analogous to transient comparator circuits in engineering, which produce a directional output based on input difference but do not retain memory of past states. This mechanism is in contrast to winner-take-all (WTA) or bistable attractor circuits, which are formed by strong local recurrent excitation and produce hysteresis. In contrast to many other *Drosophila* brain regions, where strongly recurrent excitation is commonly presented, strong recurrent excitation is not discovered in the pro-goal circuit. This is crucial. Otherwise, the system would easily overshoot when the system reaches the goal direction (*Δθ* = 0°) due to hysteresis. Biologically, the pro-goal circuit resembles Rac1-RhoA polarity systems in migrating cells ^51,52^, where mutual inhibition between signaling modules amplifies slight input differences into robust front-rear polarization.

An intriguing feature of the circuit architecture is that PFL3 neurons project to descending neurons through both a direct and an indirect pathway. The indirect route, mediated via the pro-goal circuit in the LAL, is functionally dominant: silencing this pathway abolishes goal-directed turning, whereas the direct PFL3→DN connection alone is insufficient to drive behavioral output. This raises the question: why retain a weak direct pathway if the indirect pathway is essential? We propose that the direct pathway may serve as a **priming mechanism**, bringing the descending neurons closer to threshold and thereby enabling a faster and more efficient response once the stronger signal arrives via the indirect route. This architecture mirrors feedforward excitation seen in other sensorimotor systems, where early subthreshold input facilitates rapid triggering by later-arriving signals.

Equally important, the indirect pathway provides a critical site for **modulation and behavioral flexibility**. In complex environments, multiple brain systems may compete to steer behavior based on internal goals, sensory threats, or learned contingencies. If PFL3 neurons projected solely to descending neurons, it would be difficult for other circuits, e.g. the anti-goal pathway, to inhibit or override the goal-directed response without broadly suppressing all motor output. The pro-goal circuit thus functions as a “gateable” relay, allowing other circuits to selectively modulate or suppress goal-driven turning without blocking the descending pathway entirely. This separation of command (PFL3) and control (LAL pro-goal circuit) confers a layered architecture in which behavioral responses are not only fast and directionally precise but also highly adaptable to changing motivational and environmental contexts.

### Anti-goal circuit: Override control with feedback constraint

In contrast, the anti-goal circuit receives rapid visual input via the ventral IPS pathway and promotes the turning away from the visual stimuli. However, its architecture suggests that it is not a strong independent controller. Rather, it employs three complementary mechanisms: (1) a direct but push-pull output to DNs, (2) recruitment and reversal of the pro-goal circuit, effectively **hijacking** its output to the opposite direction, and (3) an “internal” brake system formed by a feedback inhibition loop that constraints the anti-goal circuit’s own output amplitude.

This architecture mirrors the override systems in robotics and digital control, where a secondary controller cannot act alone but can reverse the polarity of the main controller. The internal brake in the anti-goal circuit is conceptually similar to interlock circuits or current limiters that ensure safety by capping output strength. In molecular biology, this resembles systems such as p53-MDM2 loop ^53,54^, where a stress response controller (p53) can override normal cell-cycle progression but is tightly regulated via feedback inhibition.

Another notable feature of the anti-goal circuit is the asymmetric interaction between the pro-goal and anti-goal pathways. While the pro-goal circuit does not inhibit the anti-goal circuit, the anti-goal pathway can actively suppress and invert the pro-goal circuit. This asymmetry suggests a functional hierarchy: when both pathways are engaged, the system prioritizes the anti-goal response. In ecological terms, this makes adaptive sense: responding to an immediate threat or aversive stimulus is typically more urgent than continuing to pursue an internal goal. The anti-goal circuit’s ability to override the pro-goal drive reflects a built-in bias toward survival-driven behavioral interruption, ensuring that ongoing goal-directed actions can be rapidly halted and reversed when necessary.

It is also important to distinguish the anti-goal response described here from a more general avoidance behavior. The anti-goal circuit does not operate in isolation; it requires the presence of a goal signal from the pro-goal pathway in order to function. In the absence of goal-related input, the anti-goal circuit cannot trigger any response, highlighting its role not as a standalone avoidance mechanism, but as a goal-modulated override system that redirects ongoing goal-directed actions away from an aversive stimulus. In contrast, a pure avoidance response should be able to trigger turning behavior independently of internal goals. Indeed, our connectome analysis reveals that the IPS projects not only to the LAL but also directly to descending neurons, potentially forming a parallel, faster-acting avoidance pathway. This architecture implies a layered defensive system: a fast, goal-independent reflex pathway for urgent avoidance, and a slower, context-sensitive anti-goal pathway that modifies goal pursuit only when necessary. This separation allows the nervous system to flexibly arbitrate between reflexive escape, goal continuation, and goal redirection, depending on internal state and environmental context.

### Symmetry breaking in behavioral stalemate

We further examined two behavioral conditions, Rear Stalemate and Front Stalemate, that challenge directional decision-making. In both cases, we observed that the push-pull architecture of the pro-goal and anti-goal circuit plays a crucial role in resolving symmetry.

The symmetry-breaking behavior resembles the metastable states in digital electronics, which linger in uncertain states until random fluctuation resolves them. The behavior is also analogous to the lateral inhibition-based patterning in developmental biology. A direct parallel is found in the Notch-Delta signaling system, where two neighboring cells with equal inputs eventually break symmetry through mutual inhibition and stochastic tipping, resulting in distinct fates ^55–57^.

### Conceptual implications: Shared principle across scales and systems

Taken together, our findings highlight how *Drosophila* LAL implements a modular control architecture that balances goal pursuit with environmental responsiveness. The underlying motifs – symmetry-sensitive push-pull comparators, feedback-limited overrides, and noise-driven decision gates -- recur across biological and engineering systems. These shared patterns suggest that evolution and engineering converge on similar solutions when facing similar problems. What distinguishes the LAL is not merely its ability to compute directional outputs, but its contextual-sensitive integration: the anti-goal circuit only becomes effective in the presence of the goal. This logic mirrors biological systems where signals are contingent on combinatorial context, which is a principle deeply embedded in gene regulation, cell signaling, and decision-making networks across levels of organization.

By tracing these analogies, we argue that the LAL does not merely encode the motor signals, but implements a compact, dynamics conflict-resolution engine, which is an insect-scale example of how compact circuits can resolve symmetry, arbitrate competing drives, and generate adaptive behaviors. These findings may inspire future efforts to build bio-inspired, neuromorphic control systems that mimic these principles to navigate complex environments with minimal hardware.

### Limitations and Future Directions

As with any modeling study, there are important limitations that should be addressed in future work. First, our circuit model assumes that all neurons operate under a simplified and uniform dynamical regime, using sigmoid activation functions to approximate their input–output relationships. While this abstraction captures key network-level computations, real neurons in the LAL may exhibit more complex intrinsic dynamics — such as oscillatory firing, spontaneous activity, or adaptation — which could influence the circuit’s temporal behavior and decision thresholds. Second, although our model already includes most of the highly connected neurons between PFL2/3 and DNs, other neurons may also participate in the pro-goal or anti-goal steering control. For example, AUTO19 or LAL083 were suggested to be part of the central complex steering neurons ^40^. Moreover, some neurons may convey information from the mushroom body for the olfaction-driven steering control ^40^. Third, our synaptic polarity assignments are based on neurotransmitter predictions from the FlyWire dataset ^58^, which were generated using machine learning classifier ^24,58^. While our assumption that GABA and glutamate are inhibitory aligns with existing Drosophila literature^59^, it remains possible that prediction errors or context-dependent transmitter actions could alter the functional logic of the circuit. Moreover, we treated the inhibitory GABA and glutamate as functionally equivalent, but in reality, they may differ in kinetics, receptor mechanisms, or downstream effects. Incorporating more biologically detailed neuron models and experimentally validated neurotransmitter identities will be essential for refining and validating the proposed circuit mechanisms.

## Methods and Materials

### The connectome data and neuron selection

The identity of neurons, connections between neurons, and the connection weights of the proposed model were retrieved from Flywire^24^, an electrical microscopy-based *Drosophila* connectome database. We assume that the connection weight between two neurons is proportional to the number of synapses between them.

We also checked the hemibrain^60^ and found that the connections between the neurons and the relative weights are consistent with the Flywire database. However, most neurons in the proposed model cross the midline, whereas hemibrain provides data for neurons in one hemisphere, except for those in the central complex. Therefore, the proposed model was built primarily based on data from Flywire.

In the present study, we investigate the role of the lateral accessory lobe (LAL) in goal-directed turning behavior (Fig. 1A). Therefore, we focused on LAL neurons that mediate signal transmission between head- and goal-direction encoding neurons and descending motor neurons. Out of more than 2212 neurons innervating each side of LAL, we performed selection criteria to focus our analysis on a small but essential subset of neurons, which includes 12 PFL2 neurons, 24 PFL3 neurons, 22 LAL neurons, and 8 DNa neurons. See Results for details.

### The neuron model

The LAL model was built and simulations were performed using the Brian2 simulator^61^. The input sources of the LAL model were the head-direction module, goal-direction module (for the pro-goal circuit), and IPS (for the anti-goal circuit). Considering that these input sources are outside the focus of this study, we constructed simplified circuits for the head- and goal-direction modules, aiming to reproduce the experimentally observed goal- and head-direction signals, rather than building biophysically realistic models. Both modules consist of an input and an output layer (Supplementary Fig. 23; Supplementary Tables 3-6), which project to the PFL2 and PFL3 neurons (Supplementary Tables 7-10). See Supplementary Methods for detailed descriptions of the two modules. Driven by the head- and goal-direction modules, the PFL2 and PFL3 neurons exhibited heading-goal offset (Δ*θ*) dependent activities that resembled observations reported previously^35,36^ (Supplementary Fig. 24).

The firing rates *r* of all neurons (except for head-, goal-direction modules and PFL2/3, which are described in Supplementary Methods) were modeled by:

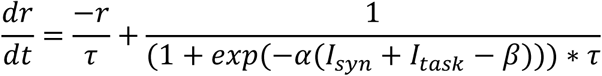

where *τ*=10ms is the time constant of the neuron, *α* and *β* are parameters for the sigmoidal activation function, *I*_*sys*_ is the synaptic current from other neurons, *I*_*task*_ is task protocol related inputs from outside the LAL circuits.

The synaptic current *I*_*syn*_ for each neuron is determined by:

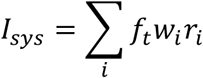

where *r*_*i*_ is the firing rate of the i-th input neuron and *w*_*i*_ is the associated synaptic weight, which is positive for an excitatory input and negative for an inhibitory input. The synaptic weights were determined based on the Flywire database. For each pair of connected neurons, we calculated the mean synapse number on the left and right sides of the brain and used this mean value in the model, assuming that the circuits were generally symmetric. The variable *f*_*t*_ is a factor dependent on the type of neurotransmitter *t*. All the model neurons are predicted (by the Flywire database) to be one of the three neurotransmitter types: cholinergic (Ach) (PFL3, DE1, DE2, DE3, VE1, VE2 and all the DNs), GABAergic (GABA_A_) neurons (DI1, and VI2) and Glutaminergic (Glut) (DI2, DI3, DI4, and VI1)(Supplementary Table 2). While cholinergic and GABAergic receptors are known to be excitatory and inhibitory, respectively, we assumed that glutaminergic neurons are inhibitory in the present study based on previous studies on *Drosophila* neurotransmitters^24,58^.

### The model tuning

The proposed model was highly constrained by the conntecome data. Only three parameters were tuned during model development. The first is the neurotransmitter factor *f*_*t*_, which was determined through parameter sweeping. The optimal values (ACh = 0.2, GABA = 0.09, Glutamate = 0.04) were those maximizing the left-right firing rate difference of the descending neurons DNa02 at 90° head-goal offset in the open-loop pro-goal task. The other two tuning parameters are *α* and *β*, which define the shape of an activation function. The parameter *β* defines the input level that drives the neuron to half of its maximum firing rate, while *α* defines the slope at this point. The parameters *α* and *β* were determined based on a homeostatic hypothesis: a neuron would adjust its response so that its firing rate (*r* = 0∼1) roughly matches the dynamical range of the input (from no input to maximum excitatory input). Based on this assumption, we set the *β* of a neuron roughly equal to half of its maximum excitatory input. But for the sake of model simplicity, which means reducing the number of distinct parameter values, we did not set each *β* to the exact value stated above, but to the closest value from the following representative numbers: 18, 20, 50, and 120. See Table 2 for details. The parameter *α* was set to equal to 5/*β*, or αβ = 5. These settings resulted in *r* ≈ 0.99 at its maximum excitatory input, and *r* ≈ 0.5 at its half excitatory input. The weight of each synapse was computed as the mean of the left and right synapse counts to ensure the symmetry of the circuit. Gaussian noise was applied to PFL3 and PFL2 populations, and the IPS input in the anti-goal conditions. The noise was equal to 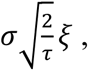 where *σ* control noise strength, *τ* = 10 ms, *ξ* was Gaussian white noise generated using Brian2.

**Table 2.** The parameters for each neuron model.

| Neuron | $\alpha$ | $\beta$ | Neuron | $\alpha$ | $\beta$ |
| --- | --- | --- | --- | --- | --- |
| DE1 | 0.25 | 18 | VI2 | 0.25 | 20 |
| DE2 | 0.25 | 18 | VE1 | 0.25 | 20 |
| DE3 | 0.25 | 18 | VE2 | 0.25 | 20 |
| DI1 | 0.1 | 50 | DNa01 | 0.25 | 20 |
| DI2 | 0.05 | 120 | DNa02 | 0.1 | 50 |
| DI3 | 0.25 | 20 | DNa03 | 0.1 | 50 |
| DI4 | 0.25 | 20 | DNa04 | 0.1 | 50 |
| VI1 | 0.25 | 20 | DNg13 | 0.25 | 20 |

### The closed-loop system

In the closed-loop system, the output of the model changed the head direction, which was fed back to the circuit model in the closed-loop experiment. Specifically, in each timestep of the simulation, the instantaneous turning velocity is determined by *Δr*_*DNa*02_ (right-left DNa02 firing rate difference) and a new head direction at the end of the timestep is computed based on the velocity. The new head direction is fed back to the neural circuit to generate new PFL2 and PFL3 input and subsequently produce a new *Δr*_*DNa*02_ for the next timestep.

The angular velocity of the body turning is computed by:

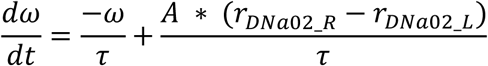

where ω is the angular velocity, *τ*=20ms, A = 0.15 is the gain that adjusts the relation between angular velocity and descending neuron activity, and *r*_*DNa*02_*R*_ and *r*_*DNa*02_*L*_ represent the firing rate of right and left DNa02 neurons, respectively.

## Data availability

This is a theoretical study based on Flywire and hemibrain, which are publicly available connectome databases.

## Code availability

The code used in the present study will be made publicly available after the manuscript is published in a peer-reviewed journal.

## Acknowledgements

C.-C. Liao thanks Dr. Ching-Che Charng for insightful discussions. C.-C. Lo thanks Dr. Ann-Shyn Chiang for advice and discussions. This work was supported by the National Science and Technology Council grants 111-2311-B-007-011-MY3, 114-2311-B-007-016-MY3, and 114-2321-B-002-027, and by the Brain Research Center under the Higher Education Sprout Project, co-funded by the Ministry of Education in Taiwan.

## Author Contribution

C.-C. Liao wrote the code, performed simulations and analysis, produced figures and wrote the manuscript. C. N. participated in the early phase of the study by writing the code, performing simulations and preparing figures. Y.-Sh. Liu wrote the manuscript and contributed the intellectual content through discussion. C.-C. Lo designed and supervised the study, wrote and finalized the manuscript.

