## Supplementary Figures for "Flexible Steering and Conflict Resolution: Pro-Goal/Anti-Goal Gating in *Drosophila* Lateral Accessory Lobes"

Supplementary Figure 1

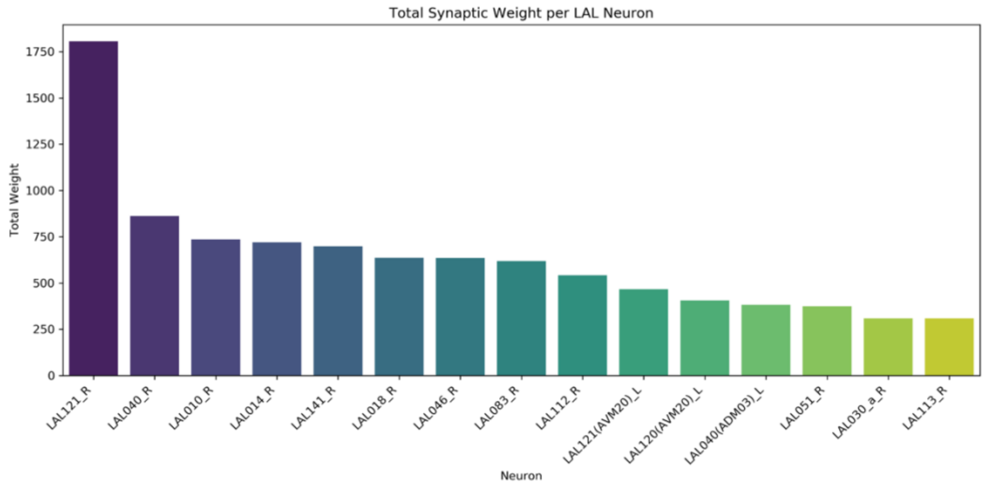


**Supplementary Fig. 1** Connection strengths of neurons that are postsynaptic to the PFL3 neurons and postsynaptic to the descending neurons (DNa01, DNa02, DNa03, DNa04, DNg13) in the lateral accessory lobe (LAL). The total weight of a neuron is defined as its synaptic counts with PFL3 and the descending neurons. The data are retrieved from NeuPrint (https://neuprint.janelia.org/).

Supplementary Figure 2

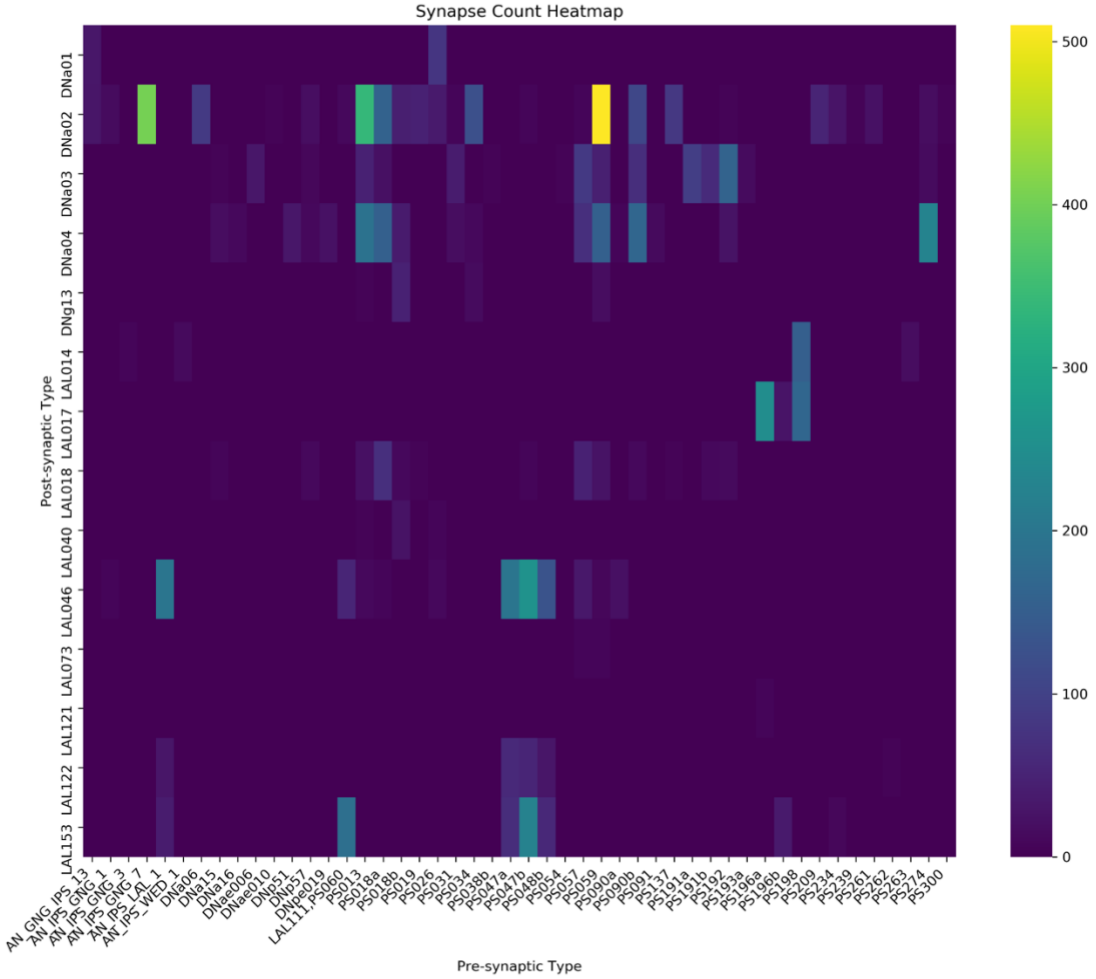


**Supplementary Fig. 2** The connectivity between IPS neurons and selected LAL neurons in our model. IPS neurons are defined as neurons with the majority of their pre or post synapses located in IPS. The data are retrieved from Flywire (https://flywire.ai/).

Supplementary Figure 3

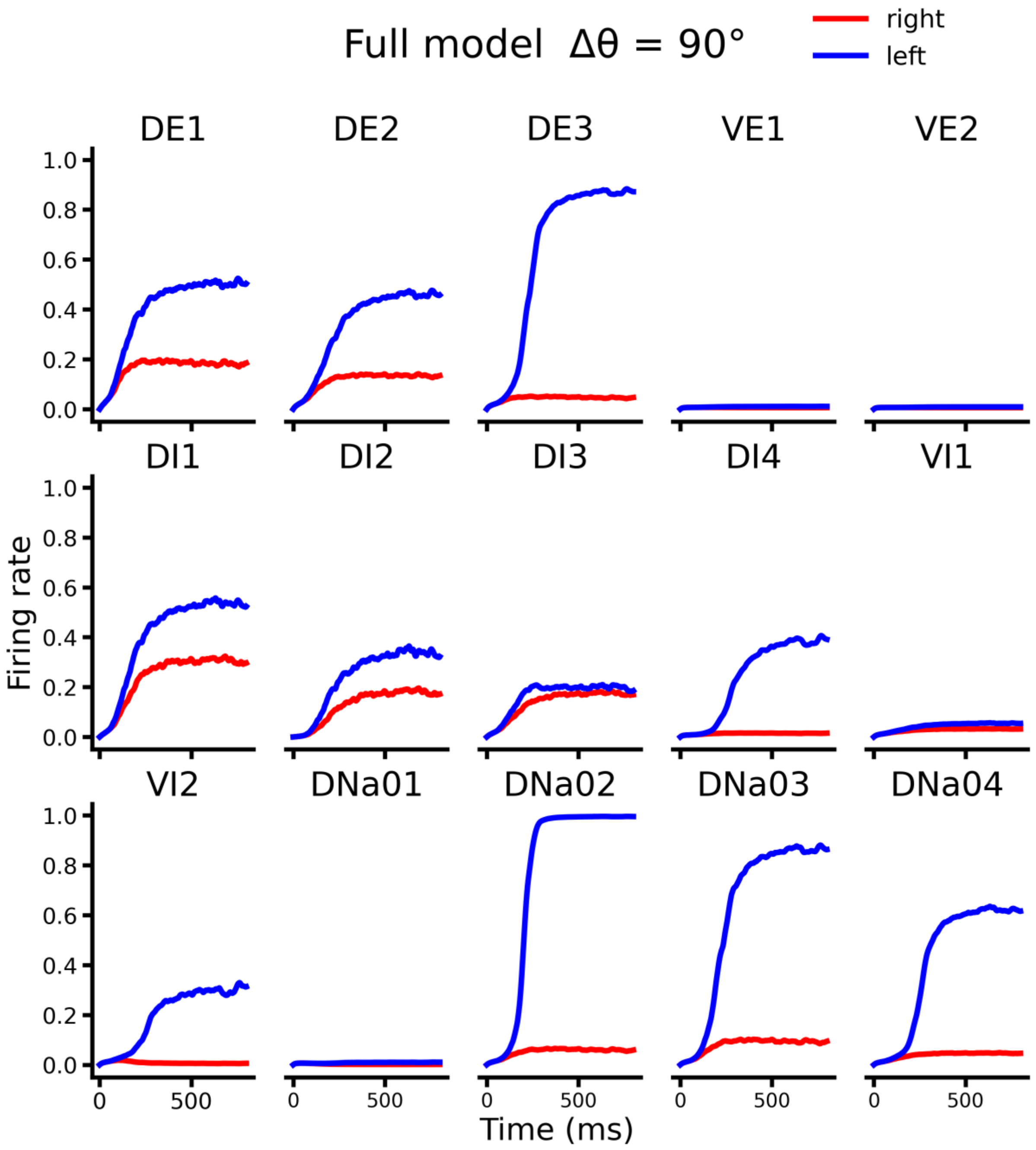


**Supplementary Fig. 3** Activity traces of all neurons in the trial shown in Fig. 2B.

Supplementary Figure 4

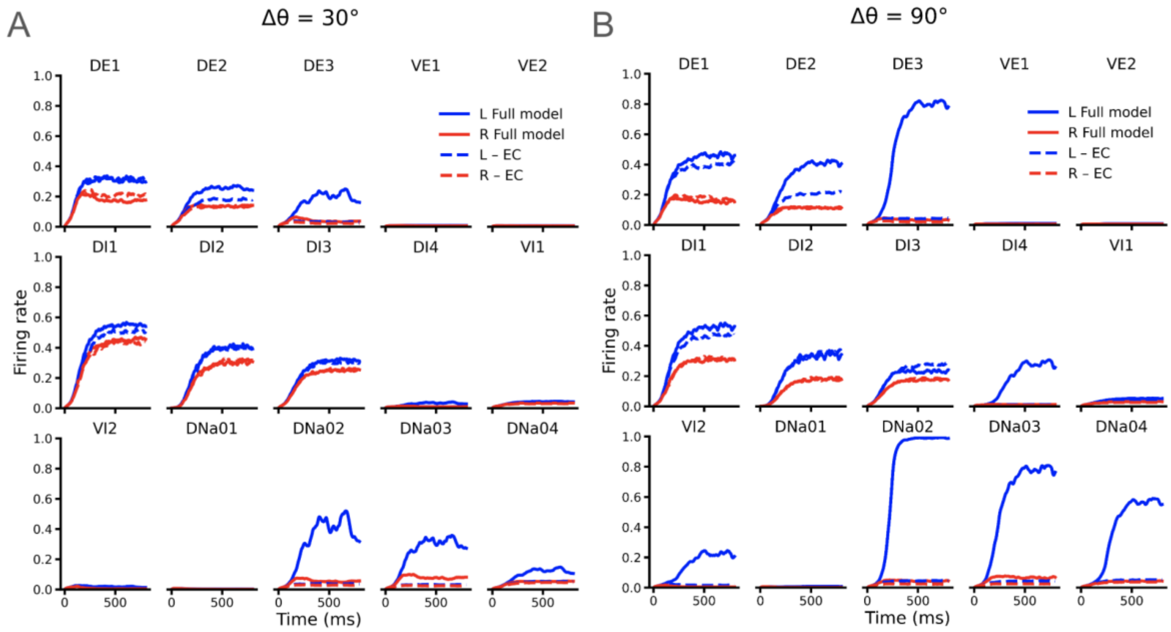


**Supplementary Fig. 4** A, B. Activity traces of all neurons in the trial shown in Fig. 2C top and bottom, respectively.

Supplementary Figure 5

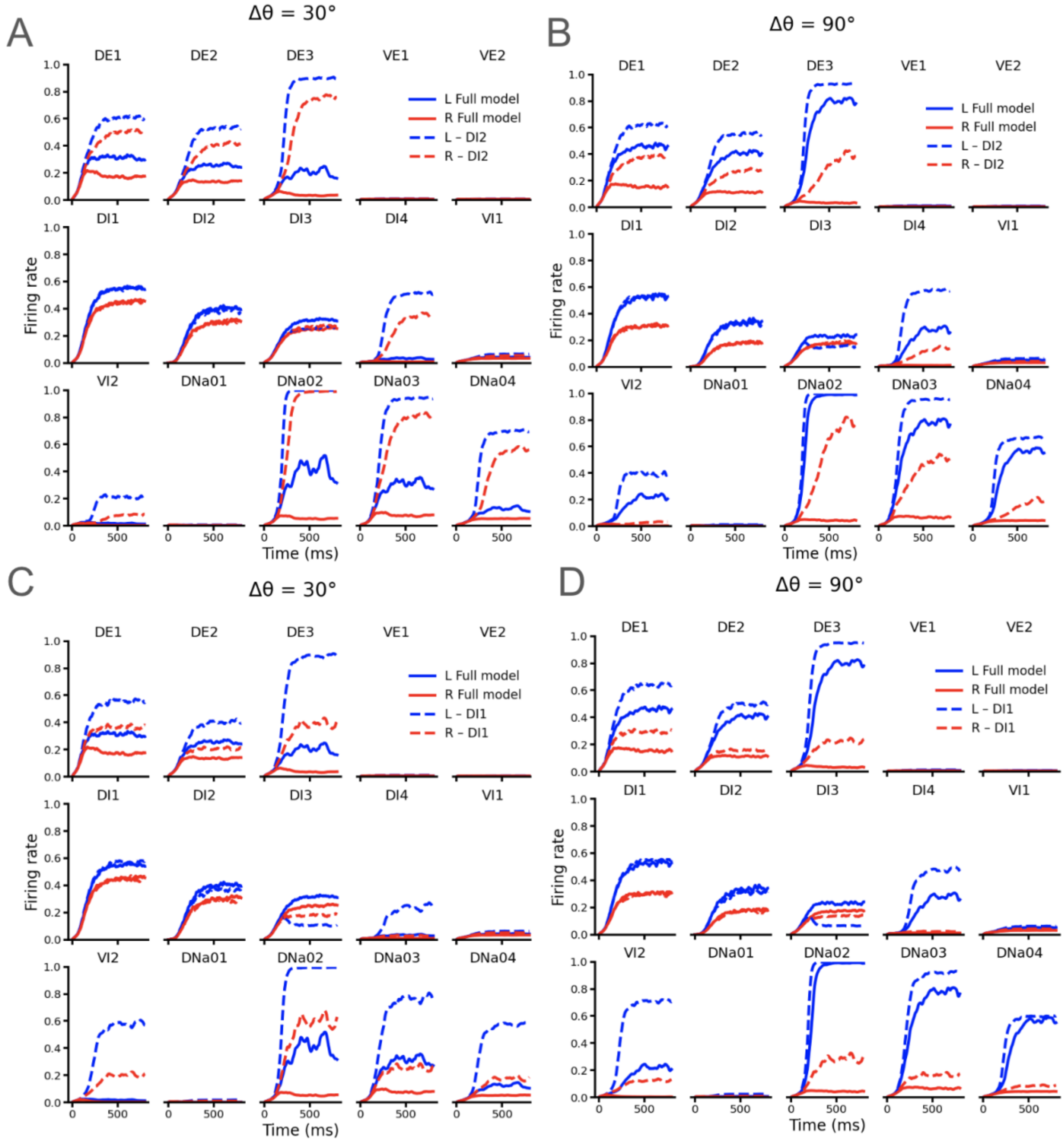


**Supplementary Fig. 5** A, B. Activity traces of all neurons in the trial shown in Fig. 2F top, bottom, respectively. C. Activity traces of all neurons in the full model and without DI1 at $\Delta\theta$=30° . D. Activity traces of individual neurons in the full model and without DI1 at $\Delta\theta$=90°.

Supplementary Figure 6

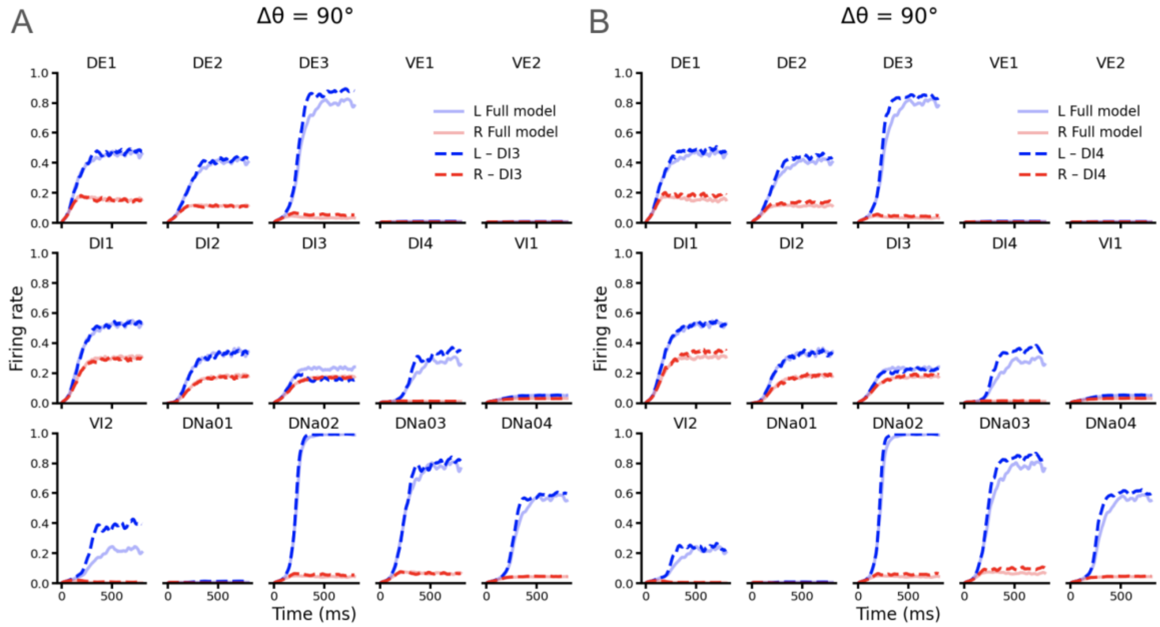


**Supplementary Fig. 6** A. Activity traces of all neurons in the full model and without DI3￼=90° . B. Activity traces of all neurons in the full model and without DI4￼=90°.

Supplementary Figure 7

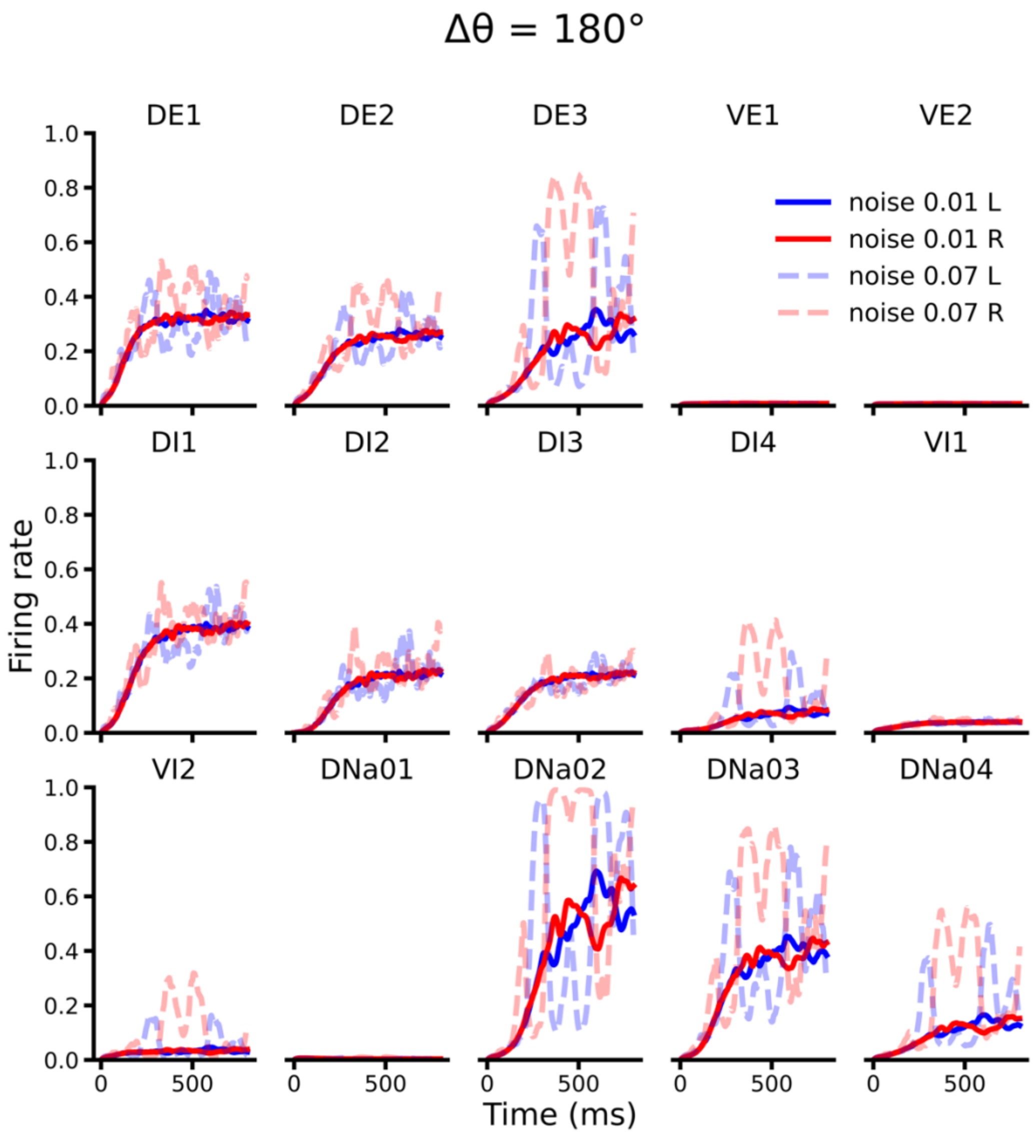


**Supplementary Fig. 7** Activity traces of all neurons in the trial shown in Fig. 3C.

Supplementary Figure 8

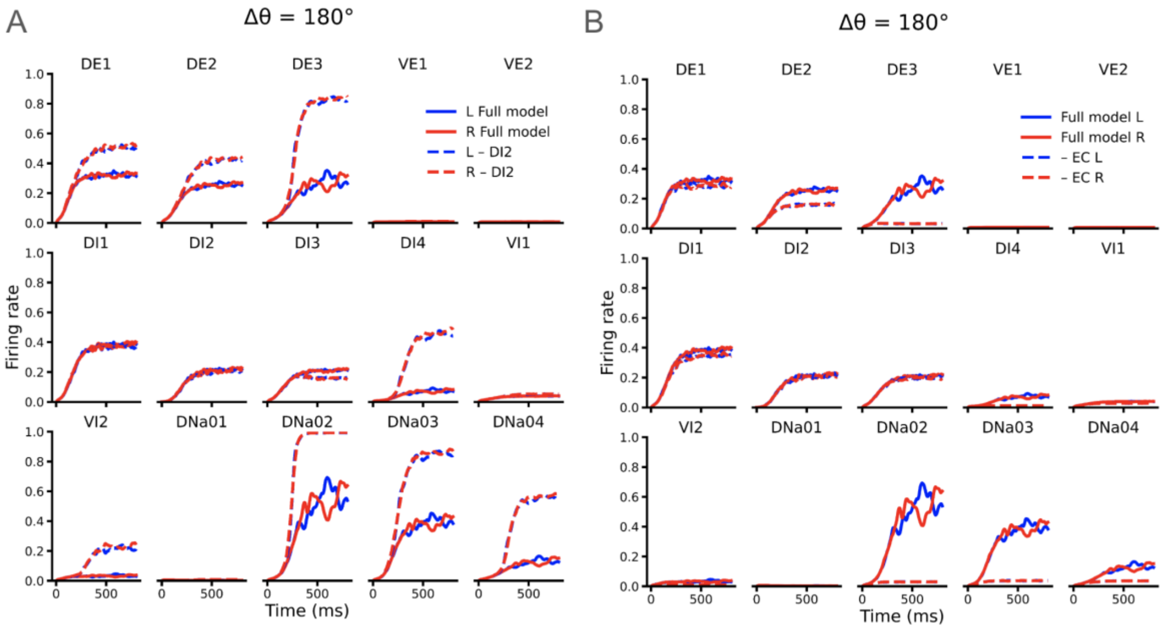


**Supplementary Fig. 8** A, B. Activity traces of all neurons in the trials shown in Fig. 3D top and bottom, respectively.

Supplementary Figure 9

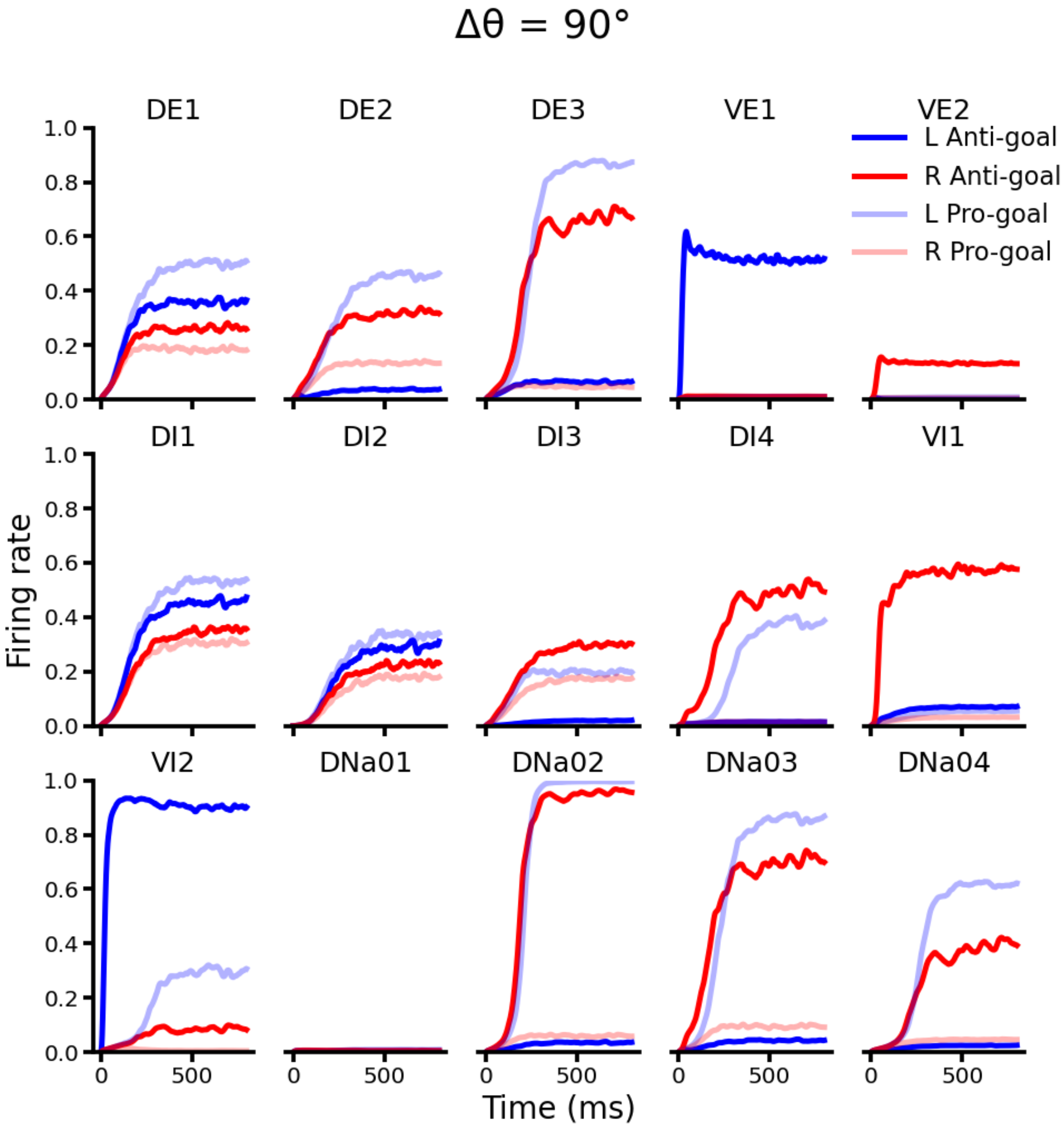


**Supplementary Fig. 9** Activity traces of all neurons in the trial shown in Fig. 4C.

Supplementary Figure 10

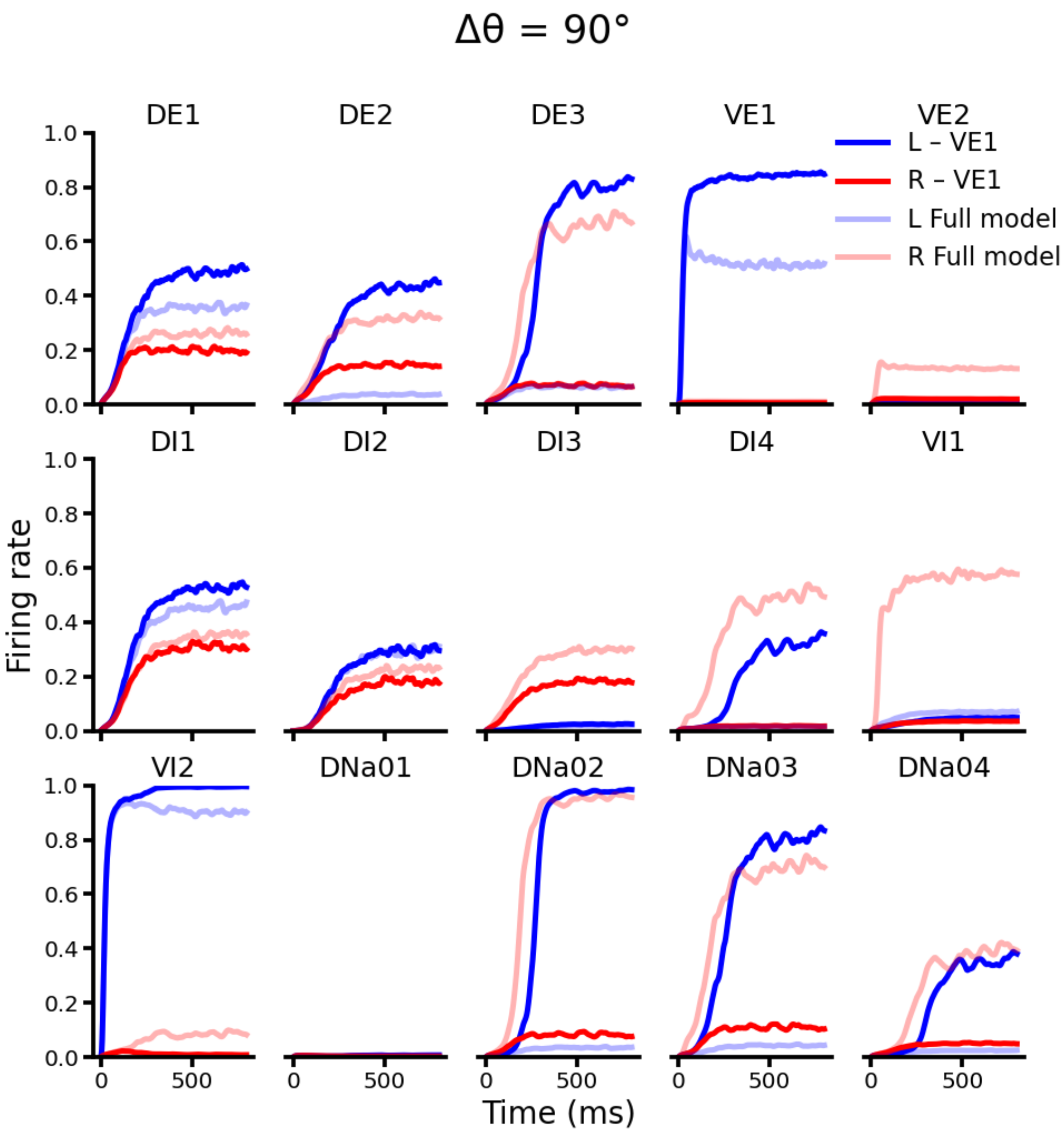


**Supplementary Fig. 10** Activity traces of all neurons in the trial shown in Fig. 4D.

Supplementary Figure 11

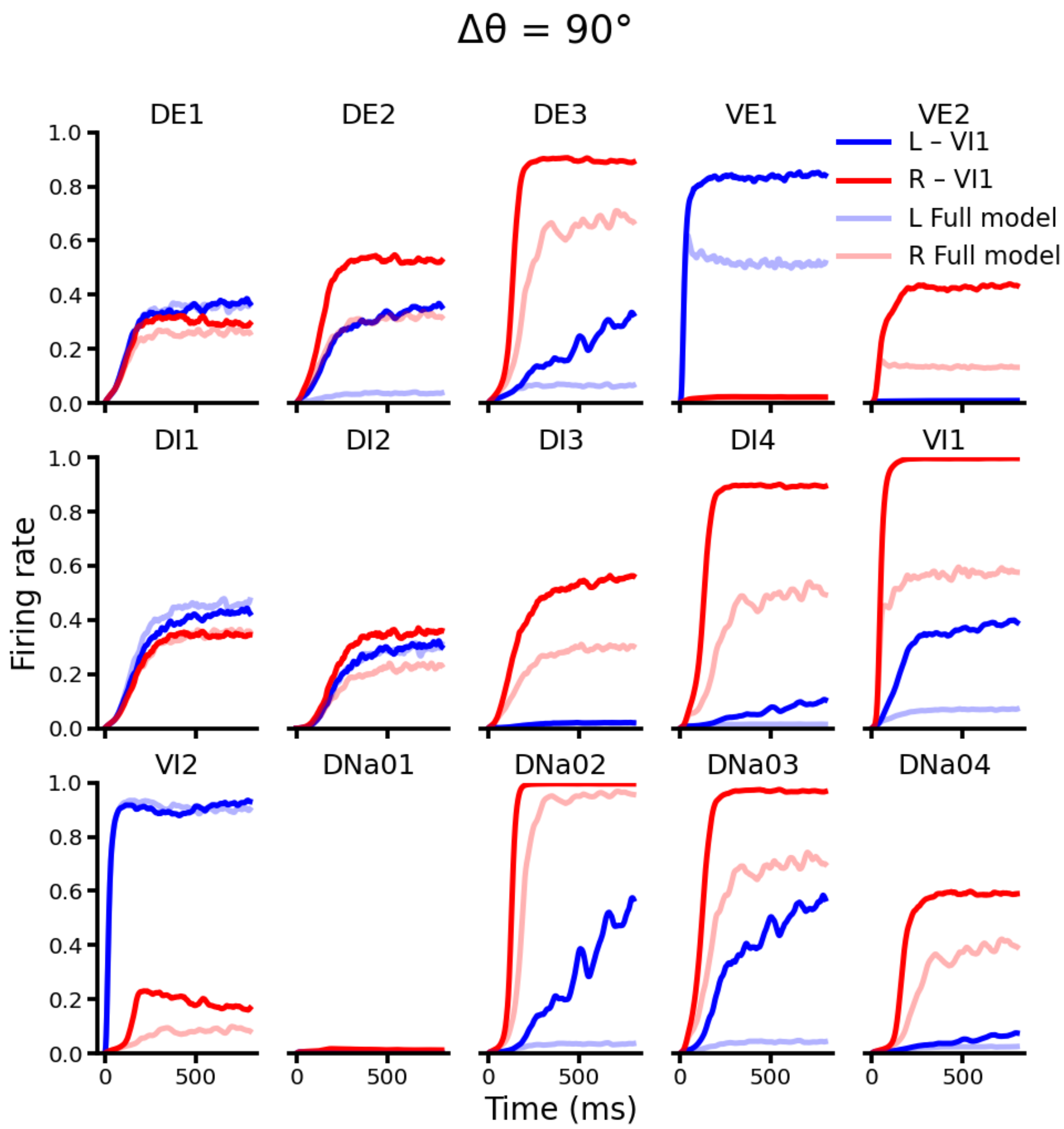


**Supplementary Fig. 11** Activity traces of all neurons in the trial shown in Fig. 4E.

Supplementary Figure 12

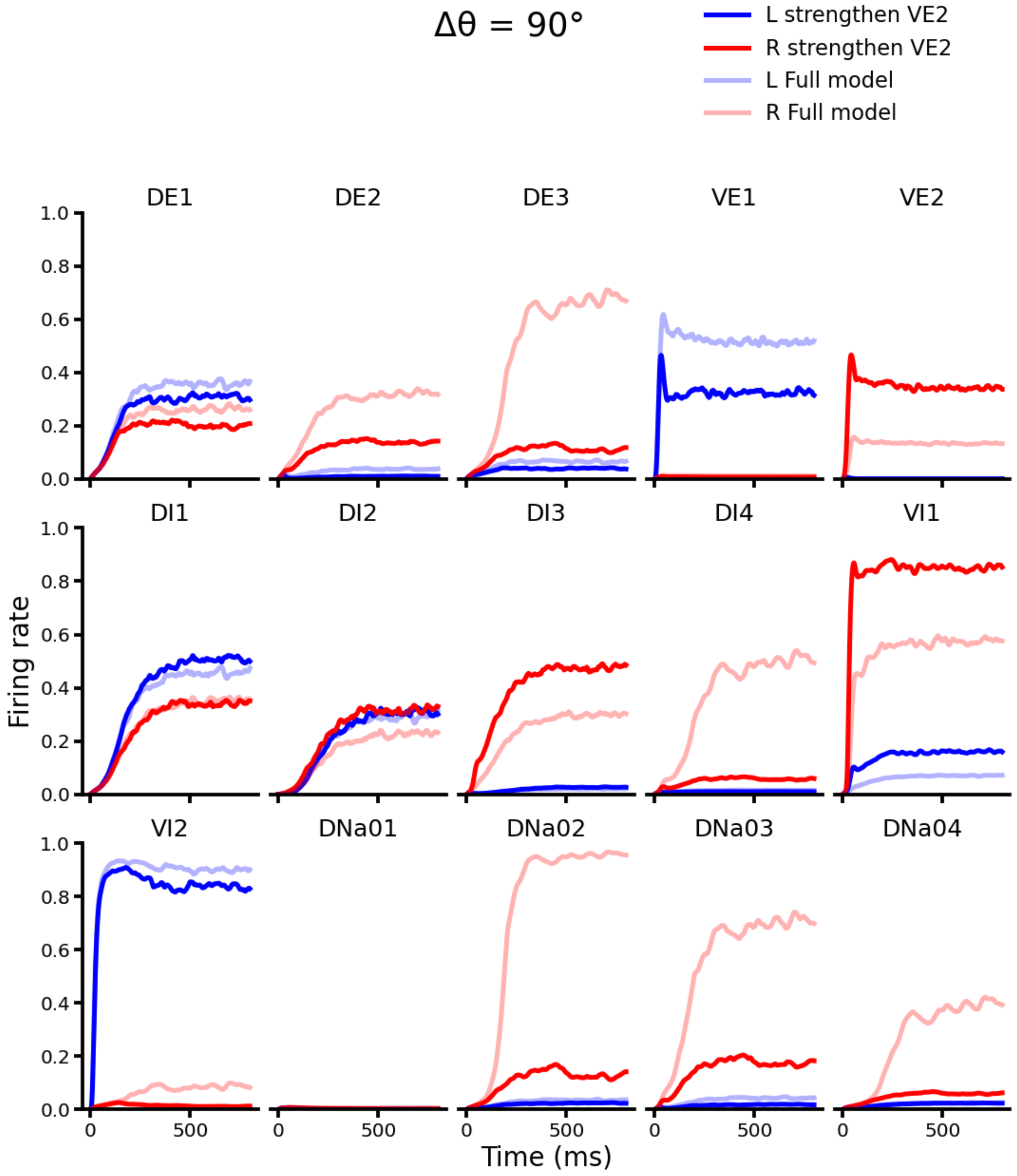


**Supplementary Fig. 12** Activity traces of all neurons in the trial shown in Fig. 4F.

Supplementary Figure 13

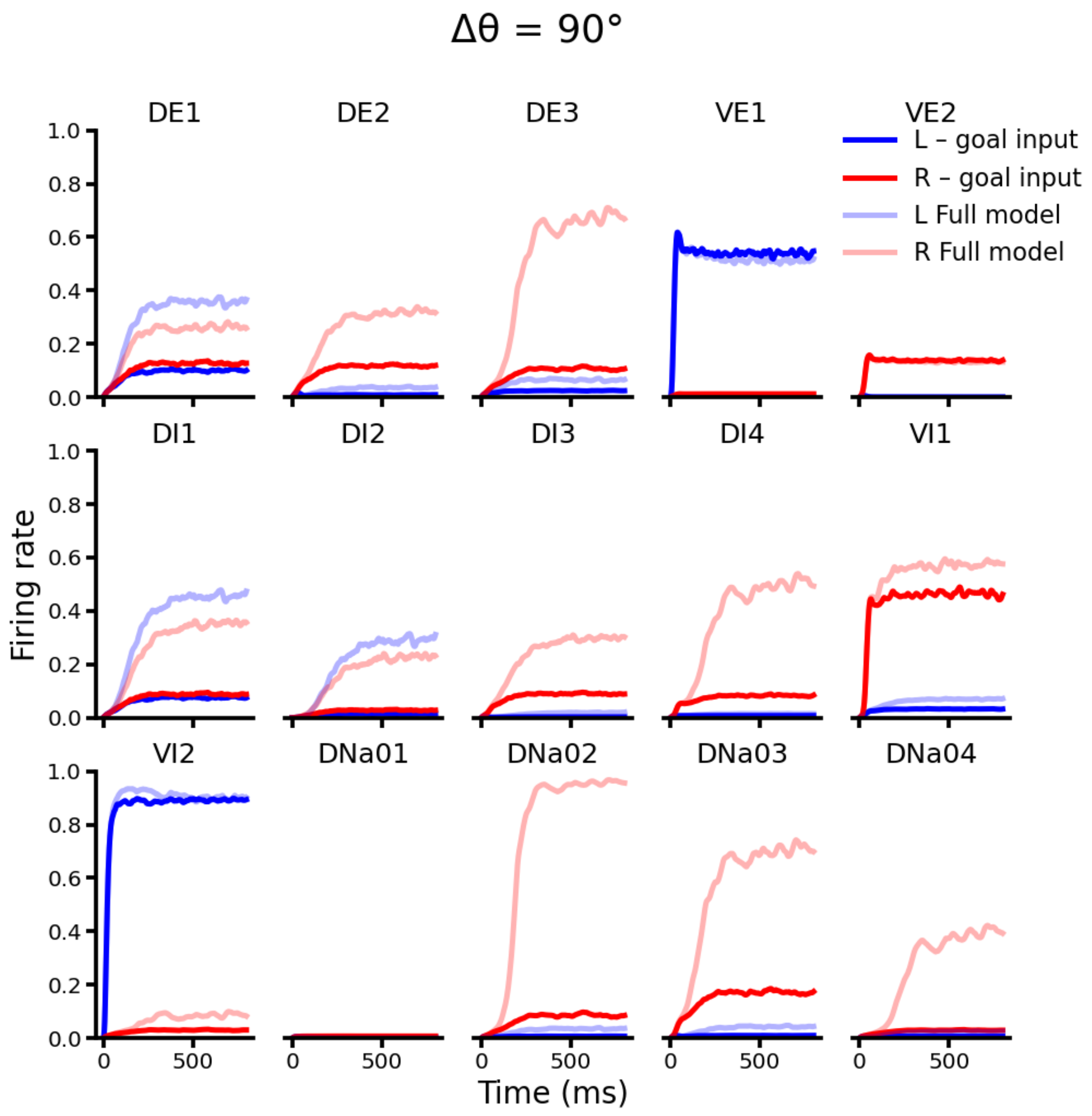


**Supplementary Fig. 13** Activity traces of all neurons in the trial shown in Fig. 4G.

Supplementary Figure 14

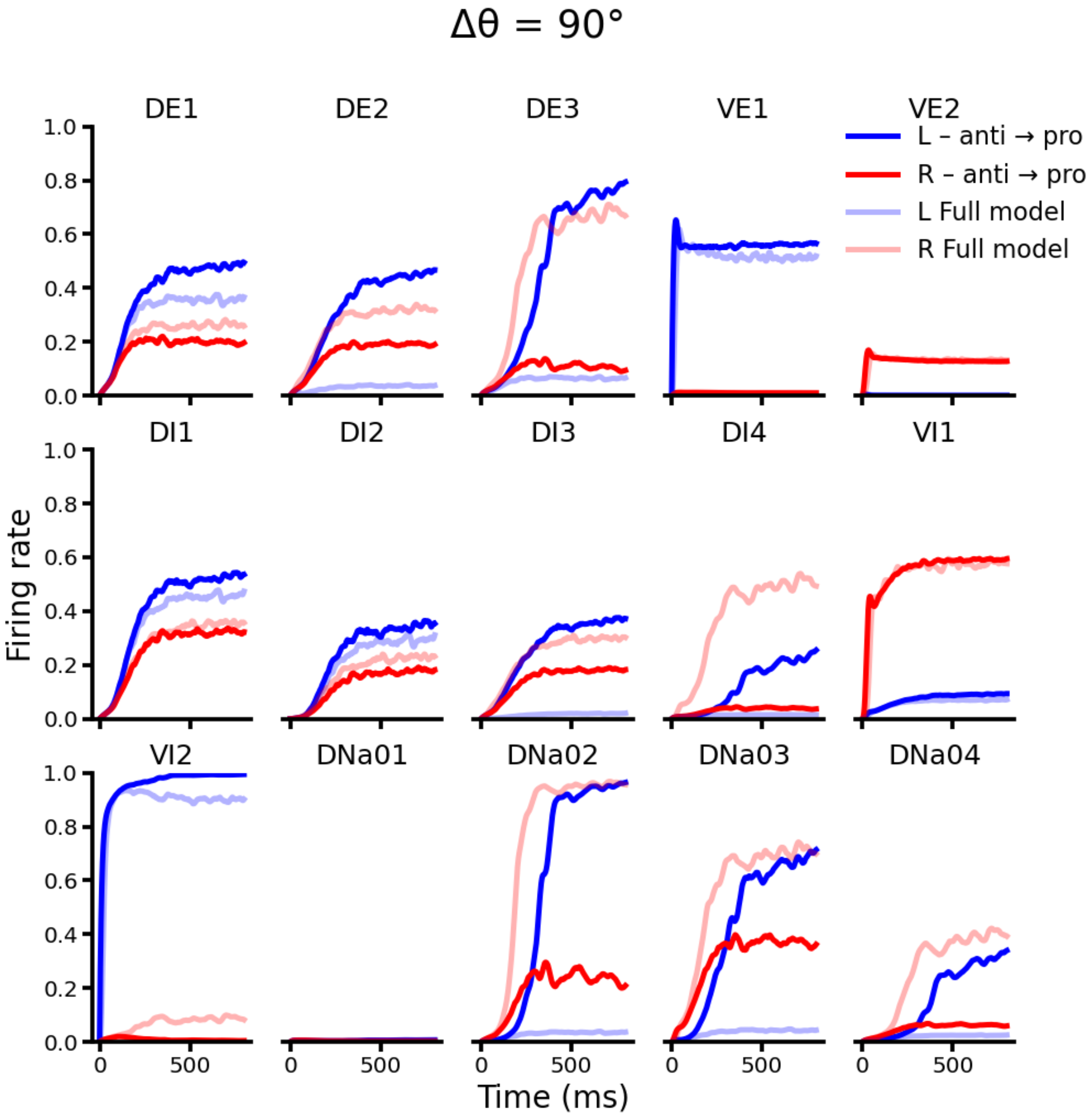


**Supplementary Fig. 14** Activity traces of all neurons in the trial shown in Fig. 4H.

Supplementary Figure 15

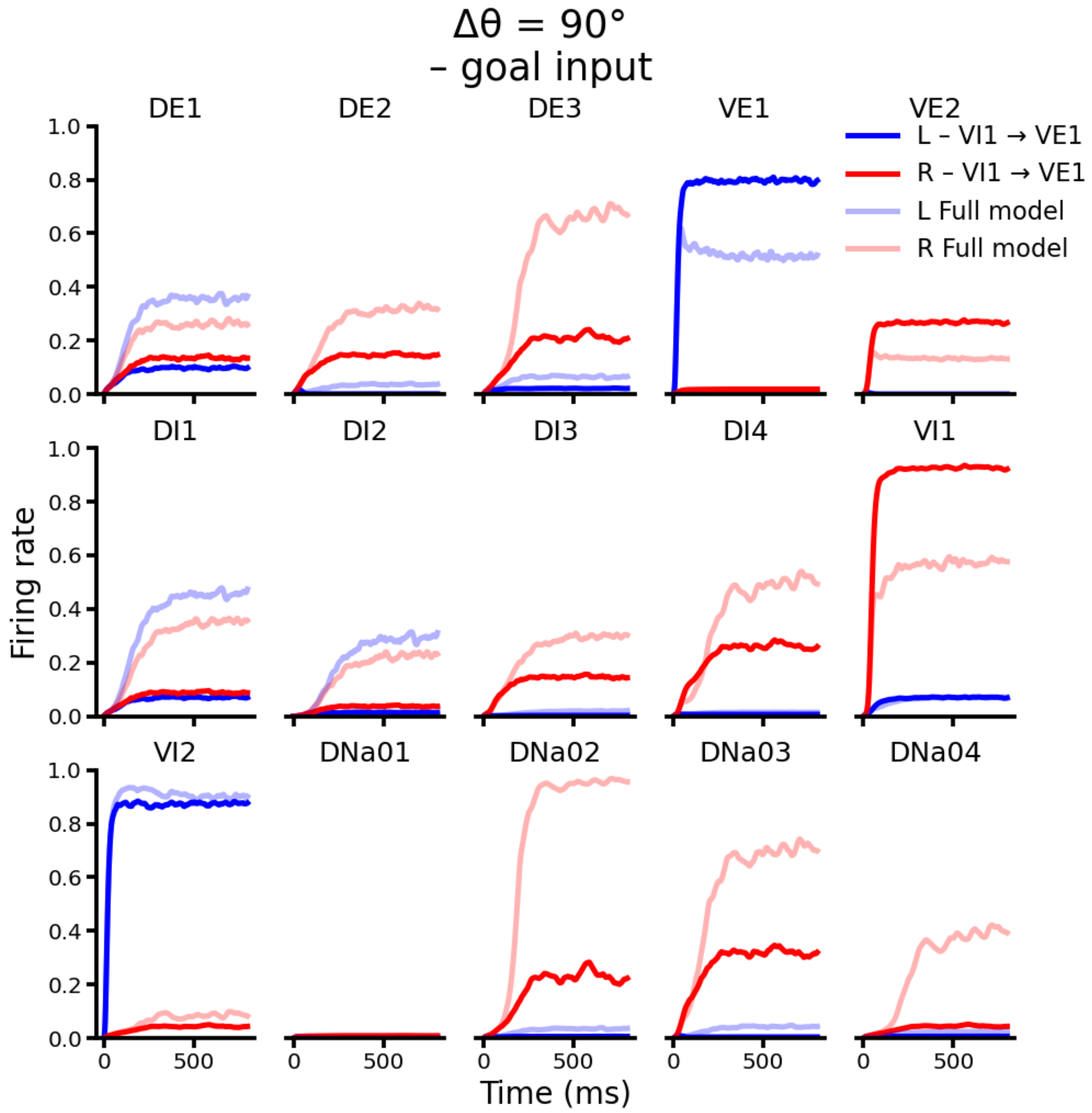


**Supplementary Fig. 15** Activity traces of all neurons in the trial shown in Fig. 4J.

Supplementary Figure 16

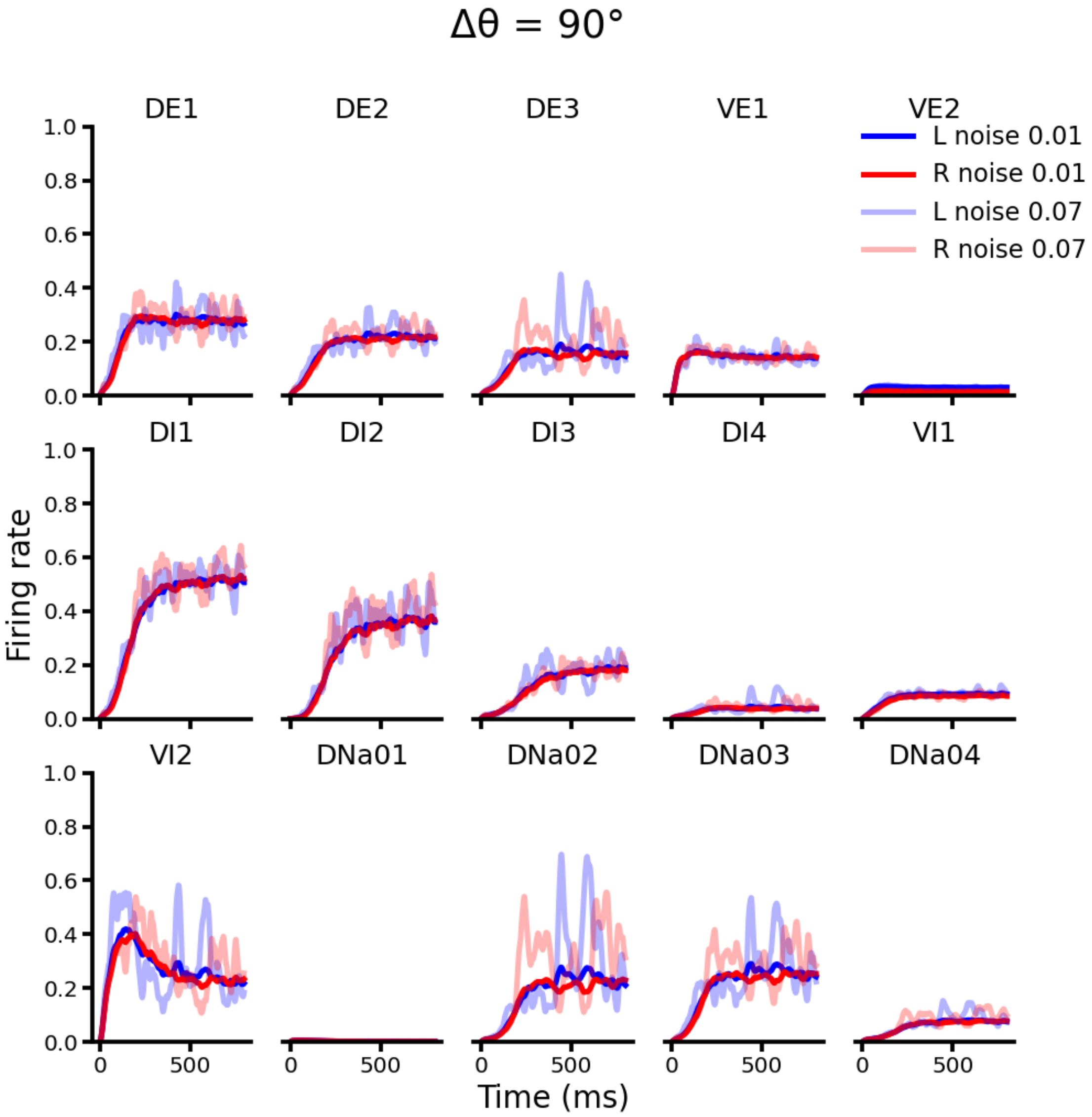


**Supplementary Fig. 16** Activity traces of all neurons in the trial shown in Fig. 4M.

Supplementary Figure 17

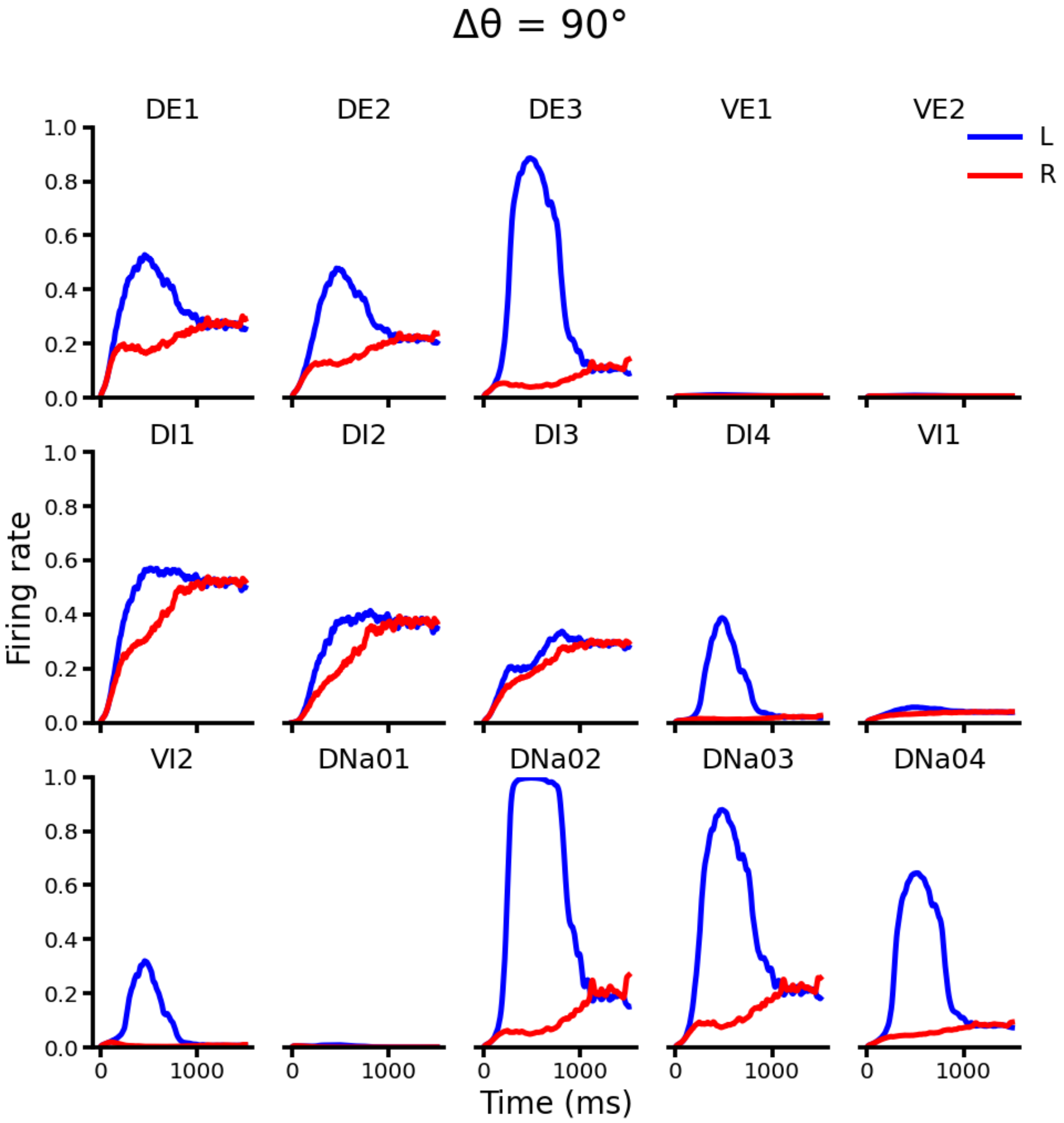


**Supplementary Fig. 17** Activity traces of all neurons in the trial shown in Fig. 5B.

Supplementary Figure 18

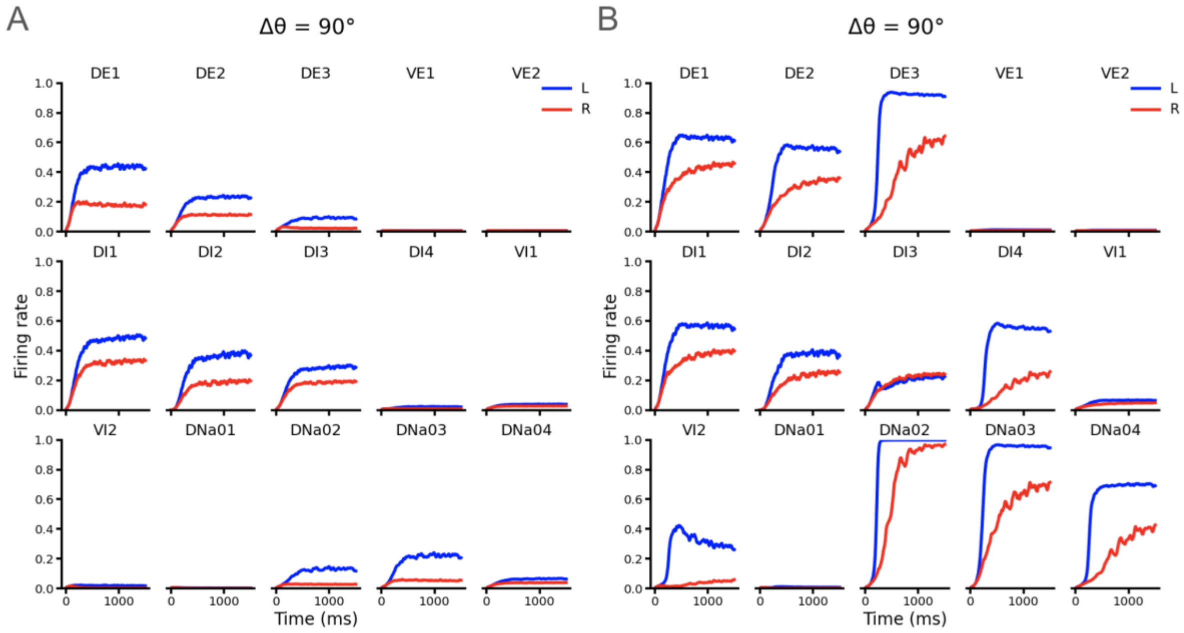


**Supplementary Fig. 18** A, B. Activity traces of all neurons in the trials shown in Fig. 5C & D, respectively.

Supplementary Figure 19

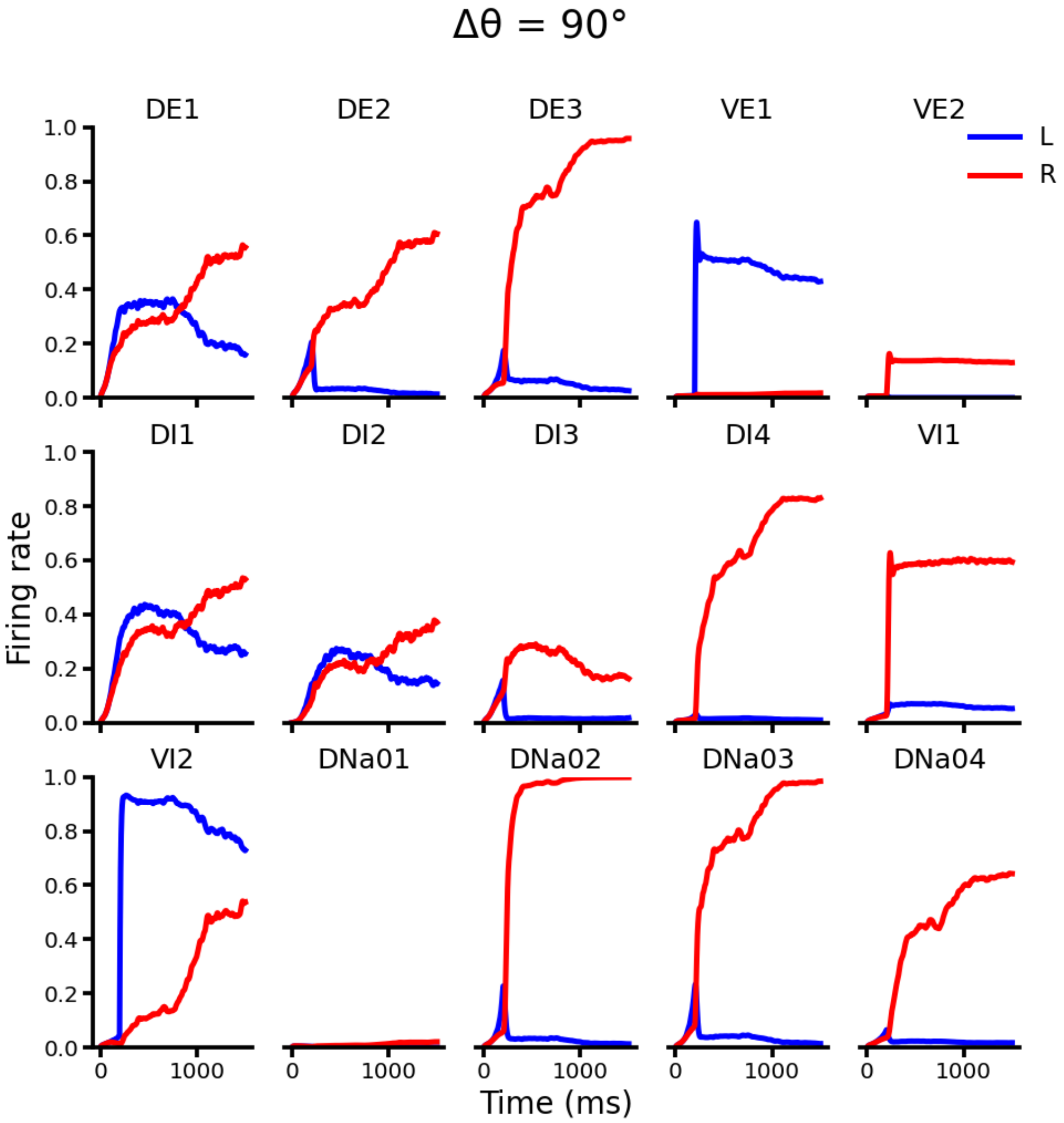


**Supplementary Fig. 19** Activity traces of all neurons in the trial shown in Fig. 5F.

Supplementary Figure 20

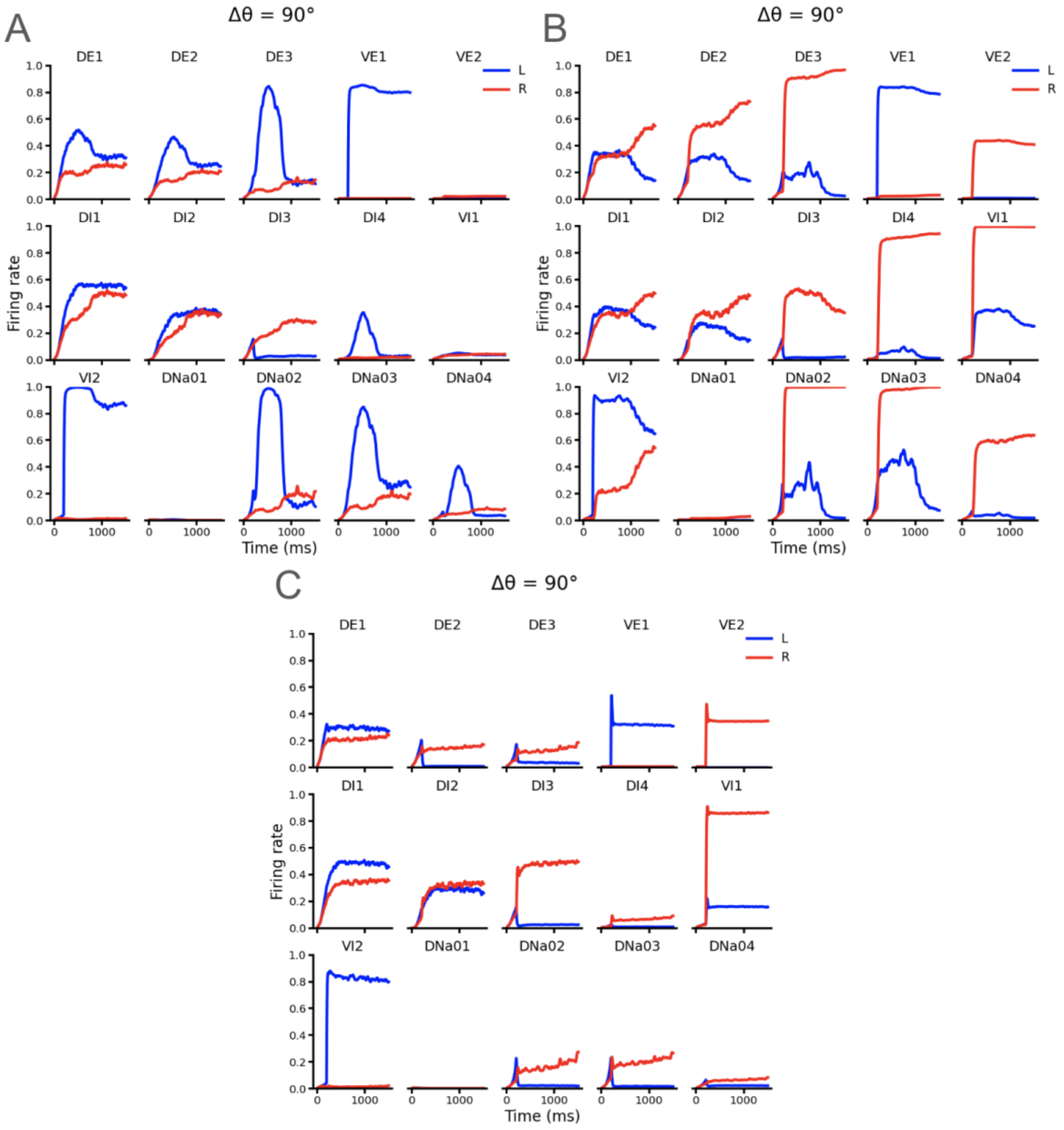


**Supplementary Fig. 20** A, B, C. Activity traces of all neurons in the trials shown in Fig. 5G left, middle and right, respectively.

Supplementary Figure 21

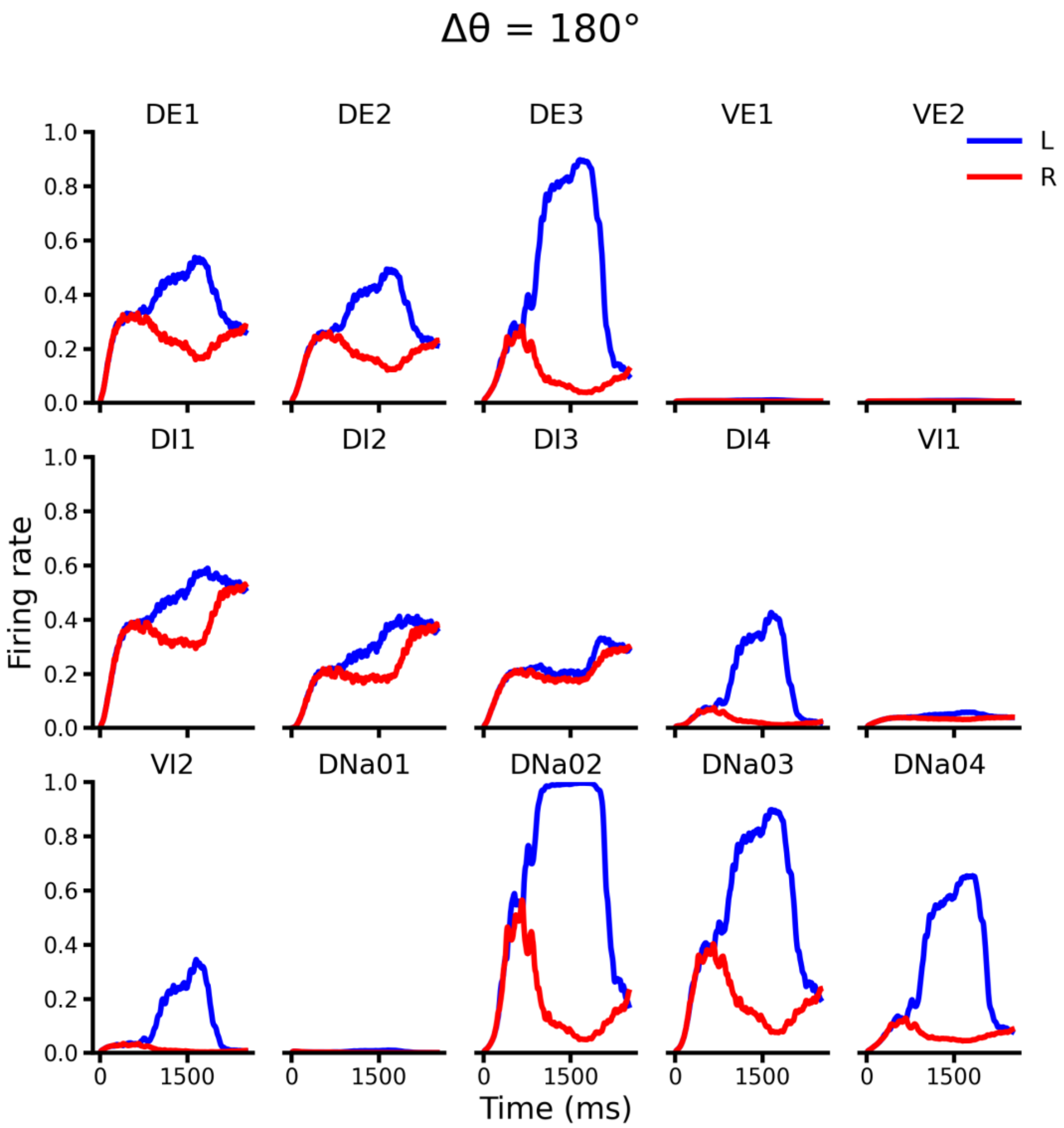


**Supplementary Fig. 21** Activity traces of all neurons in the trial shown in Fig. 6A.

Supplementary Figure 22

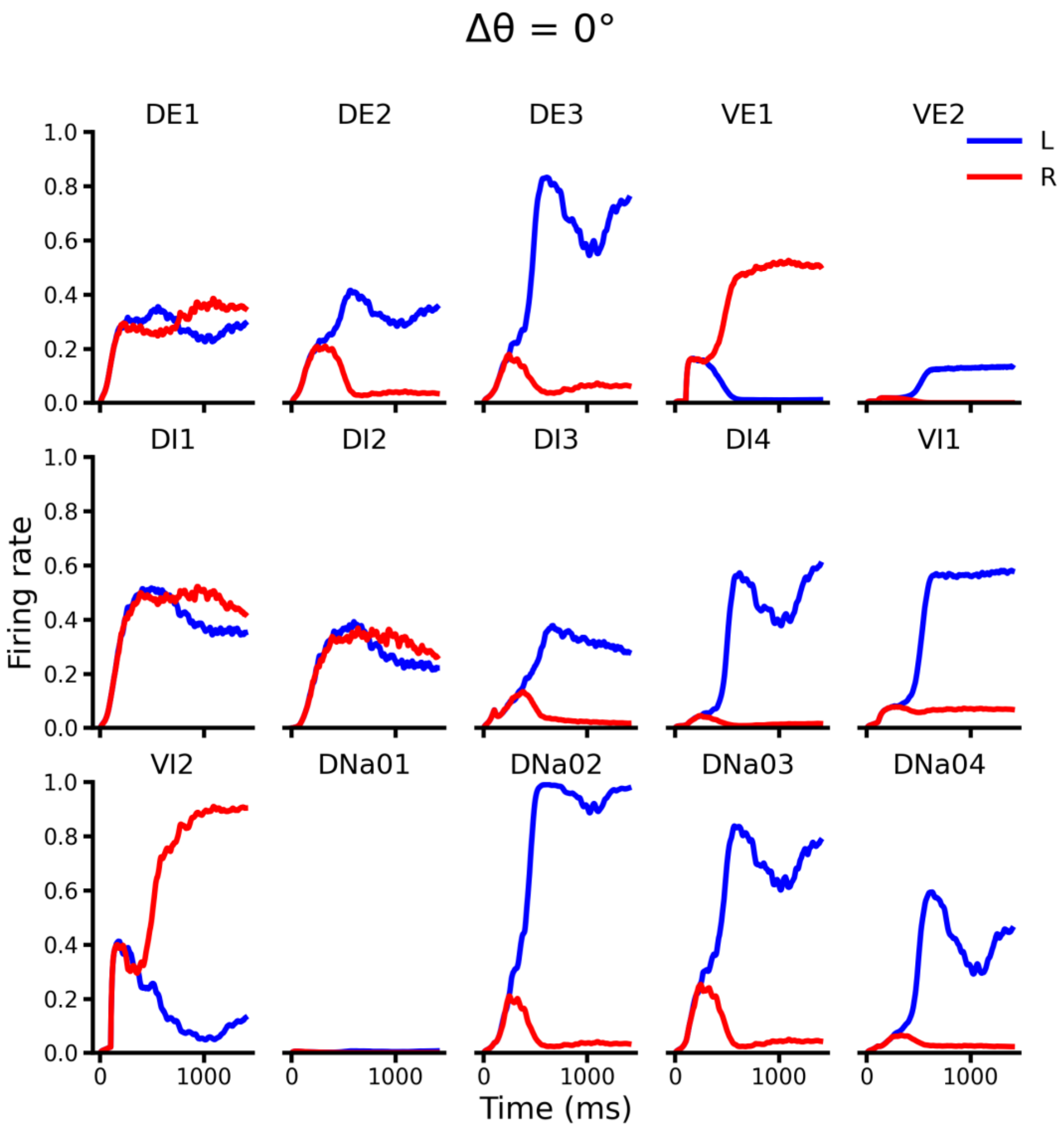


**Supplementary Fig. 22** Activity traces of all neurons in the trial shown in Fig. 6C.

Supplementary Figure 23

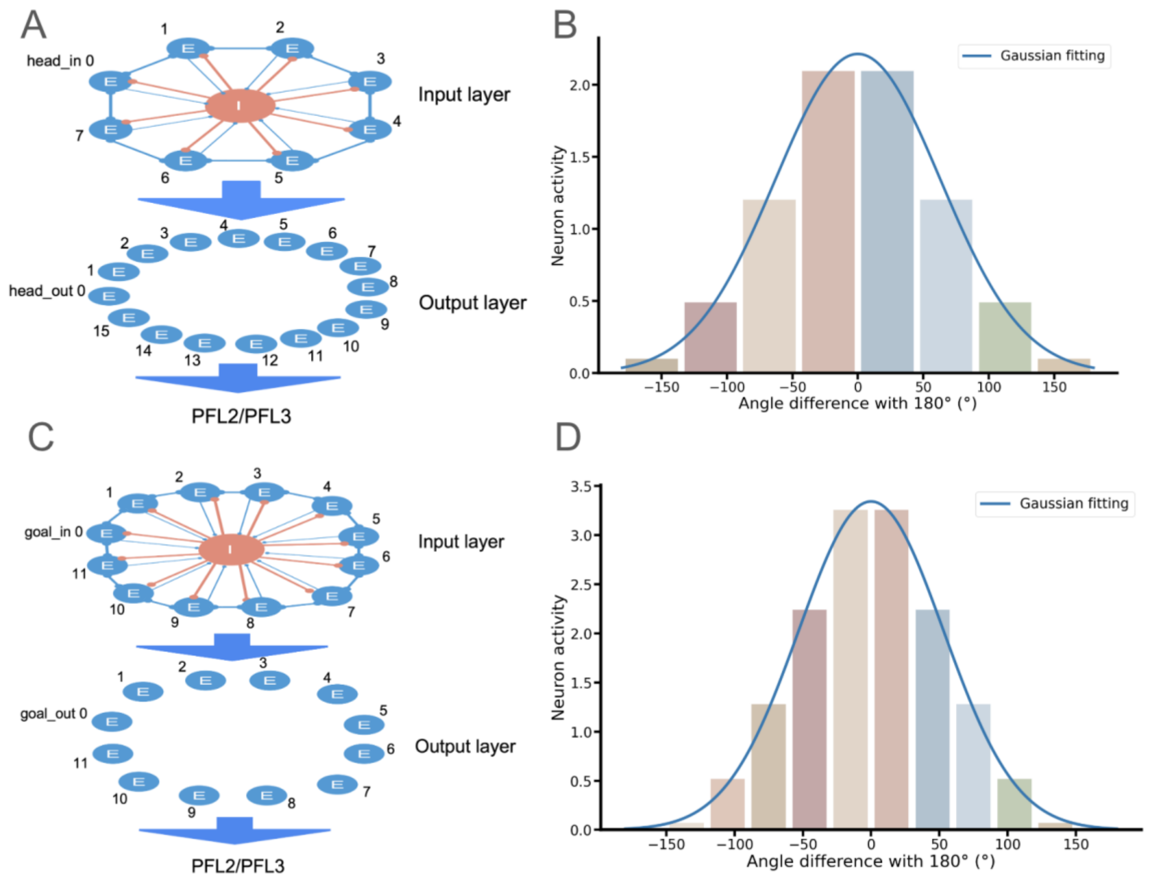


**Supplementary Fig. 23** A. Schematics of the head direction module, which consists of an input and an output layer. See Supplementary Tables 3 & 4 for the connection table and weights. B. Activities of the output layer neurons in the head direction module, exhibiting an observed Gaussian bump with a width of 89.1°. C. Schematics of the goal direction module, which consists of an input and an output layer. See Supplementary Tables 5 & 6. D. Activities of the output layer neurons in the goal direction module, exhibiting an observed Gaussian bump with width of 75.0°.

Supplementary Figure 24

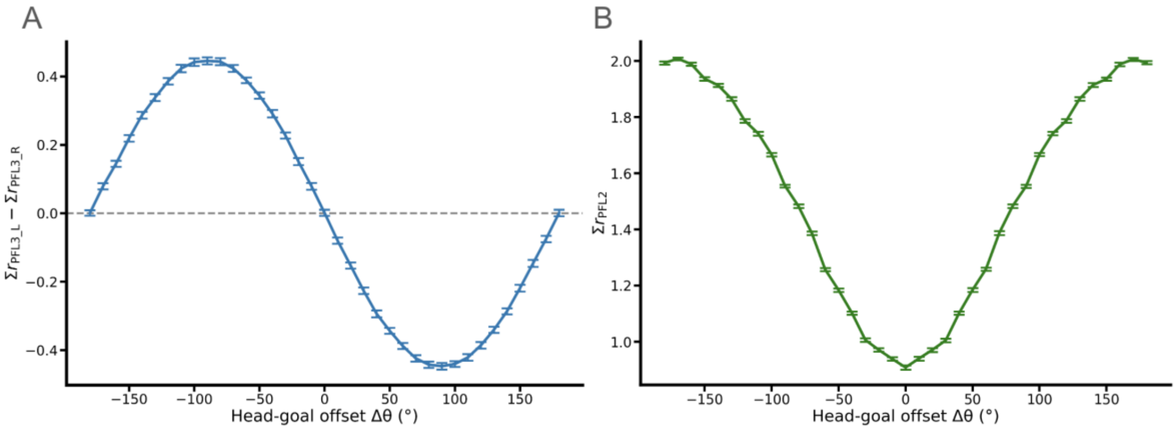


**Supplementary Fig. 24** A. PFL3 activity with noise at different head-goal offset $\Delta\theta$. B. PFL2 activity with noise at different head-goal offset $\Delta\theta$.
