## Supplementary Method for "Flexible Steering and Conflict Resolution: Pro-Goal/Anti-Goal Gating in *Drosophila* Lateral Accessory Lobes"

### Supplementary Methods

#### The head-direction and goal-direction modules

The head-direction and goal-direction modules provided input to the PFL2 and PLF3 neurons. The modules were constructed using Brin2[1]. Considering that these input sources are outside the focus of this study, we constructed simplified circuits for the head- and goal-direction modules, which consisted of an input and an output layer (Supplementary Fig. 23; Supplementary Tables 3-6). The input layer formed a ring attractor circuit with local recurrent excitation and global feedback inhibition (Supplementary Tables 3 & 5). Each neuron in the input layers encoded a preferred direction. The preferred directions were [337.5°, 22.5°, 67.5°, 112.5°, 157.5°, 202.5°, 247.5°, 292.5°] for the head-direction module, $\theta_k^{head}$ (*k*=1-8), and were [15°, 45°, 75°, 105°, 135°, 165°, 195°, 225°, 255°, 285°, 315°, 345°] for the goal-direction module, $\theta_k^{goal}$ (k=1-12). The input layers were connected to the output layers (Supplementary Tables 4 & 6), which in turn projected to the PFL2 and PFL3 neurons (Supplementary Tables 7-10).

During simulations, the activation of the input layer neurons in each module was determined by the following rules. For a given head or goal angle $\theta$, if it exactly matched an input layer neuron's preferred direction, only this neuron received an input, $I = 1$. If not, two neurons k1 and k2 with the nearest preferred directions were identified such that $\theta_{k1} < \theta < \theta_{k2}$ (mod 360∘). Then, the input was distributed between these two neurons according to a linear interpolation:

$$I_{k1} = 1 - ratio, I_{k2} = ratio$$

where

$$ratio = \frac{\theta - \theta_{k1}}{\theta_{k2} - \theta_{k1}}$$

Neurons in both modules were modeled by a simplified firing rate equation:

$$\frac{dr}{dt} = \frac{-(r + I)}{\tau}$$

where $\tau = 10ms$, and $I$ is the input current. For the input layers, $I$ is the summation of the head or goal direction activation as described above and the synaptic input ($\sum w_i r_i$, with $w_i$ and $r_i$ being the weight and firing rate of $i$-th presynaptic neuron) from other neurons in the same layer. For the output layers, $I$ is the synaptic input from the input layers. See Supplementary tables 3-6 for the connection tables and weight values for the two modules.

The PFL3 model in the present study was developed based on Mussells Pires et al. 2022[2], which suggested that the PFL3 neurons received inputs from both head-direction and goal-direction neurons. Following Mussells Pires et al. 2022[2], we modeled the firing rate $r_i$ of a PFL3 neuron $i$ by:

$$\frac{dr_i}{dt} = \frac{-r_i}{\tau} + a\log(1 + \exp(b(\sum_j w_{ij} r_j^{head} + c\sum_k g_{ik} r_k^{goal} + d))).$$

$r_j^{head}$ is the firing rate of the $j$-th output layer neuron from the head-direction module, weighted by $w_{ij}$. $r_k^{goal}$ is the firing rate of the $k$-th output layer neuron from the goal-direction module, weighted by $g_{ik}$. The parameter c (=0.67) sets the relative strength between head and goal direction signal. The rest parameters, a (=22.09 Hz), b (=2.33) and d (=-0.52), also followed the values used in Mussells Pires et al. 2022[2]. See Supplementary Tables 7 & 8 for the connection tables and weights between the output layers of both modules and PFL3 neurons.

Unlike PFL3 neurons, which innervate the contralateral LALs, PFL2 neurons innervate bilateral LALs[3–5]. The firing rate $r_i$ of the $i$-th PFL2 is given by:

$$\frac{dr_i}{dt} = \frac{-r_i}{\tau} + \frac{1}{(1+exp(-\alpha*(\sum_j w_{ij} r_j^{head} + \sum_k g_{ik} r_k^{goal} - \beta)))*\tau}$$

The weight $\alpha$ was set to 2.5, and $\beta$ was set to 2.0. Both modeled PFL2 & 3 neurons exhibited firing rate profiles identical to those described in Mussells Pires et al. 2022[2] (Supplementary Fig. 24). Supplementary Tables 9 & 10 for the connection tables and weights between the output layers of both modules and PFL2 neurons.

IPS input was given to ipsilateral LAL152 (VE1), contralateral LAL017 (VE2) and ipsilateral LAL046 (VI2). The dynamics of IPS input $I$ was described by:

$$\frac{dI}{dt} = -\frac{(I - I_{IPS})}{\tau_I} + noise$$

Where $I_{IPS}$ is the steady-state strength of the IPS input and $\tau_I$ (=10 ms) is the time constant of $I \tau_I = 10\ ms.$
